# Monospecific and bispecific monoclonal SARS-CoV-2 neutralizing antibodies that maintain potency against B.1.617

**DOI:** 10.1101/2021.12.21.473733

**Authors:** Lei Peng, Yingxia Hu, Madeleine C. Mankowski, Ping Ren, Rita E. Chen, Jin Wei, Min Zhao, Tongqing Li, Therese Tripler, Lupeng Ye, Ryan D. Chow, Zhenhao Fang, Chunxiang Wu, Matthew B. Dong, Matthew Cook, Guilin Wang, Paul Clark, Bryce Nelson, Daryl Klein, Richard Sutton, Michael S. Diamond, Craig B. Wilen, Yong Xiong, Sidi Chen

## Abstract

COVID-19 pathogen SARS-CoV-2 has infected hundreds of millions and caused over 5 million deaths to date. Although multiple vaccines are available, breakthrough infections occur especially by emerging variants. Effective therapeutic options such as monoclonal antibodies (mAbs) are still critical. Here, we report the development, cryo-EM structures, and functional analyses of mAbs that potently neutralize SARS-CoV-2 variants of concern. By high-throughput single cell sequencing of B cells from spike receptor binding domain (RBD) immunized animals, we identified two highly potent SARS-CoV-2 neutralizing mAb clones that have single-digit nanomolar affinity and low-picomolar avidity, and generated a bispecific antibody. Lead antibodies showed strong inhibitory activity against historical SARS-CoV-2 and several emerging variants of concern. We solved several cryo-EM structures at ∼3 Å resolution of these neutralizing antibodies in complex with prefusion spike trimer ectodomain, and revealed distinct epitopes, binding patterns, and conformations. The lead clones also showed potent efficacy *in vivo* against authentic SARS-CoV-2 in both prophylactic and therapeutic settings. We also generated and characterized a humanized antibody to facilitate translation and drug development. The humanized clone also has strong potency against both the original virus and the B.1.617.2 Delta variant. These mAbs expand the repertoire of therapeutics against SARS-CoV-2 and emerging variants.

## Introduction

In the ongoing coronavirus disease 2019 (COVID-19) pandemic, severe acute respiratory syndrome coronavirus (SARS-CoV-2) has infected over 270 million individuals, resulting in more than 5 millions of deaths around the globe ^1^. Although multiple vaccines are available, breakthrough infections still occur ^2^, especially with variant strains. Thus, broadly effective therapeutic options are critical for medical treatment. Monoclonal antibodies (mAbs) have been effectively deployed for prevention or treatment of COVID-19 ^3^. However, the emergence of mutations in spike ^4^ and new variant lineages calls for developing additional therapeutic interventions, including mAbs with potent and broadly neutralizing ability.

Certain variants affect the rate of spread and/or even the ability to evade immune recognition, potentially dampening the efficacy of antibody therapy or vaccines and have been designated by WHO and CDC as “variants of concern” (VoC) ^3^. The B.1.1.7 lineage (Alpha variant) has an increased rate of transmission and higher mortality ^5^. The B.1.351 lineage (Beta variant) has an increased rate of transmission, resistance to antibody therapeutics, and reduced vaccine efficacy ^6–8^. The lineage B.1.617 including B.1.617.1 (Kappa variant), B.1.617.2 (Delta variant) and B.1.617.3 have emerged and became dominant in multiple regions in the world ^9, 10^. The B.1.617 lineage has an increased rate of transmission, shows reduced serum antibody reactivity in vaccinated individuals, and exhibits resistance to antibody therapeutics under emergency use authorization (EUA) ^11–15^. The Delta variant has become dominant in US and many countries across the globe^16–18^. A newly emerged variant Omicron (B.1.1.529) with extensive mutations in the Spike gene also shows rapid transmission^19–21^. With these VoCs, even emergency-use authorized (EUA) mAb therapies have face challenges with resistant viral variants, causing some to be withdrawn, highlighting a need for more mAb candidates in development. Discovery of neutralizing antibodies with broad neutralizing activities or multi- specific antibodies, which can increase antibody efficacy and prevent viral escape, might improve the countermeasure arsenal against COVID-19 ^3^.

The majority of pre-clinical and clinical SARS-CoV-2 mAbs were discovered utilizing the blood of COVID- 19 patients ^3^. In comparison, immunization of mice followed by hybridoma screening is a standard method for discovering of therapeutic mAbs against viruses^22^. However, hybridoma screening for potent neutralizing mAbs from immunized mice is a slow and laborious process. The recent development of high-throughput single-cell technologies enabled direct sequencing of fully recombined VDJ sequences of B cell receptor (BCR) repertoires from single cells ^23, 24^. This technology has successfully been used to isolate human neutralizing mAbs against pathogens such as HIV, Ebola viruses, and recently SARS-CoV-2 ^25, 26, 27–29^.

The SARS-CoV-2 surface spike glycoprotein (S) mediates entry into target cells and is a primary target of neutralizing antibodies ^30–32^. SARS-CoV-2 spike is a trimer in the prefusion form, consisting of three copies of S1 and S2 subunits ^33, 34^. The S1 subunit is composed of an N-terminal domain (NTD) and a C-terminal receptor binding domain (RBD) that recognizes the host angiotensin-converting enzyme 2 (ACE2) receptor on the cell surface ^31, 35–37^. The S2 subunit contains the fusion peptide, along with other key regions, to induce membrane fusion of the virus and the target cell ^38^. Spike RBD is flexible, moving between “up” and “down” conformations but only binds to ACE2 in the up conformation ^34, 36, 39^. Previous studies have suggested that most of the potent SARS-CoV-2 neutralizing antibodies target spike RBD through an interface directly overlapping with the ACE2-binding surface ^40, 41^.

Here, we report the rapid identification of highly potent SARS-CoV-2-neutralizing mAbs using high-throughput single-cell BCR sequencing from mice immunized with purified SARS-CoV-2 RBD. We generated a bispecific antibody with two antigen-recognition variable regions from two of the top mAb clones against SARS-CoV-2 RBD. These monospecific and bispecific antibodies displayed high affinity and avidity to the RBD antigen, and potently neutralized historical and B.1.617 lineage viruses. We also generated and characterized a humanized antibody to facilitate clinical development, and showed strong potency against both the original virus and the Delta variant, both *in vitro* and *in vivo*. We resolved the three-dimensional cryo-EM structures of these neutralizing antibodies in complex with the spike-ectodomain trimer, which showed the epitopes as well as the combinations of open and close RBD conformations of trimeric spike bounded to the mAbs. Structure-based mutation and epitope analysis revealed the neutralization potency of the lead mAb clones against B.1.617 variants.

## Results

### Single cell BCR sequencing of RBD-immunized mice identified enriched BCRs encoding strong mAbs against SARS-CoV-2

To generate potent and specific mAbs against SARS-CoV-2, we immunized two different mouse strains: C57BL/6J and BALB/c with RBD protein with a C-terminal hexahistidine tag following a standard 28-day immunized protocol (**Extended Data Fig. 1a**). Using anti-mouse CD138 beads, we isolated progenitor B cells and plasma B cells from spleen, lymph node, and bone marrow of selected immunized mice that showed high serum binding titers against RBD (**Extended Data Fig. 1a**). We performed single cell VDJ (scVDJ) sequencing on the isolated B cells (**Extended Data Fig. 1a**). The scVDJ data revealed the landscape of immunoglobulin clonotypes in immunized mice (**Fig. 1a; Dataset S1**), and identified enriched IgG1 clones (**Fig. 1b and Dataset S1**). We took the VDJ sequences 11 top-ranked IgG1+ clones by clonal frequency. We then cloned the paired variable region of heavy and light chain into human IgG1 (hIgG1) heavy and light chain backbone vectors separately, for antibody reconstruction utilizing the Expi293F mammalian expression system.

**Figure 1.**
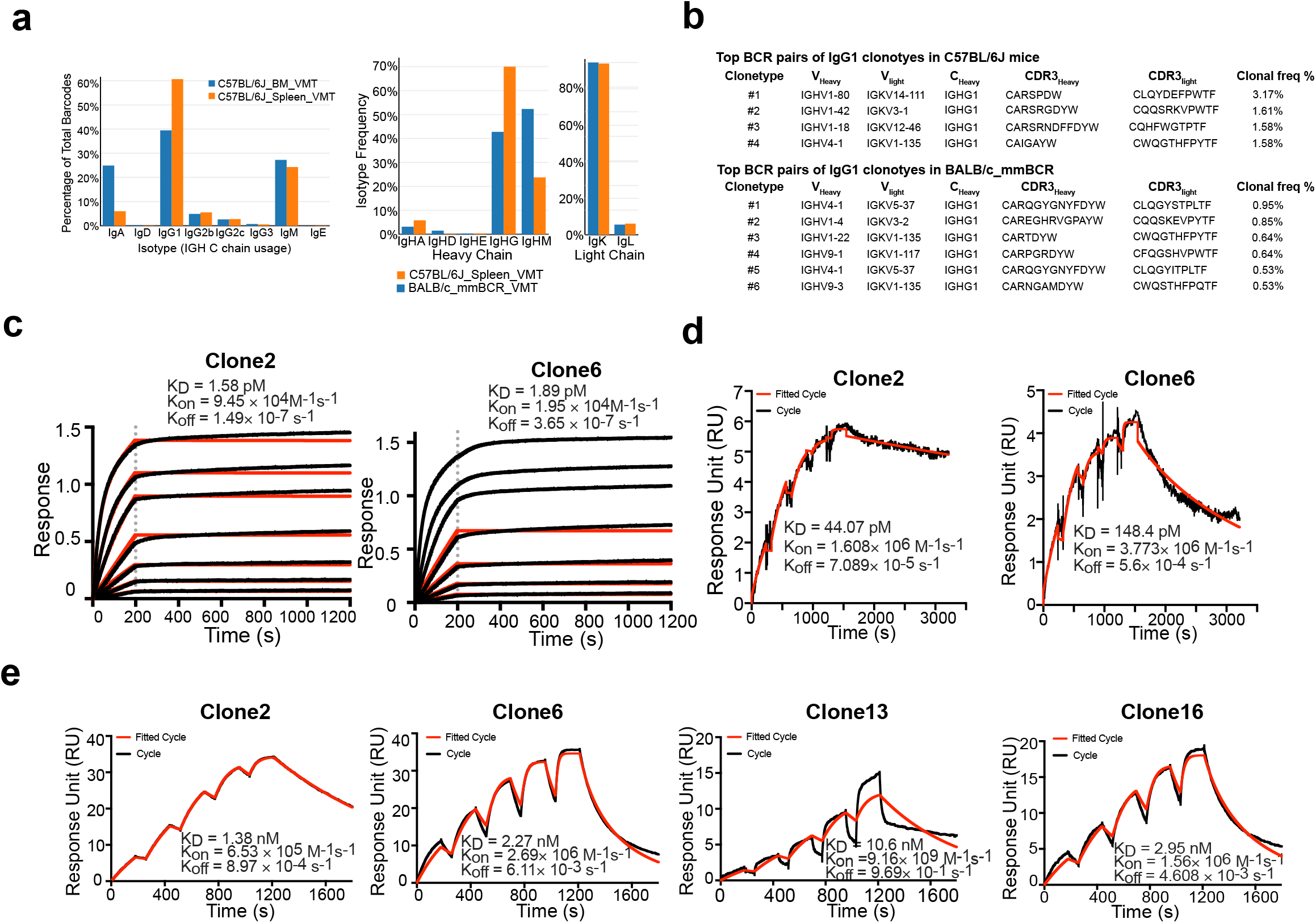
Immunization-single B cell sequencing leads to development of potent monoclonal antibodies in both monospecific and bispecific forms. **a**, Frequency distribution of immunoglobulin isotypes in SARS-CoV-2 Spike RBD immunized mice. (Top) Analysis of isotypes in both bone marrow and spleen in RBD immunized C57BL/6J mice. (Bottom) Analysis of isotypes in RBD immunized C57BL/6J and BALB/c mice. **b**, CDR3 sequences of heavy and light chains of the top enriched antibody clones from RBD-his tag immunized C57BL/6J mice (Top) and BALB/c mice (Bottom). **c**, Octet measurement of binding strengths top SARS-CoV-2 RBD-specific monospecific mAb clones (Clones 2 and 6). The binding was particularly strong, thus that the dissociation stage was never observed in this BLI assay. **d**, SPR measurement of binding strengths top SARS-CoV-2 RBD-specific monospecific mAb clones (Clones 2 and 6), using an NTA chip where the antigen (RBD) was fixed on the chip. **e**, SPR measurement of binding strengths top SARS-CoV-2 RBD-specific monospecific mAb clones (Clones 2 and 6), a humanized mAb clone (Clone 13), and a bispecific mAb (Clone 16), using a Fc chip where antibodies were fixed on the chip. Source data and additional statistics for experiments are provided in a supplemental excel file.

### Anti-Spike RBD monoclonal antibodies have single-digit nanomolar affinity and low-picomolar avidity

After expression and purification of hIgG1 antibody clones, we tested their reactivity against SARS-CoV-2 spike RBD by ELISA. Eight of the eleven mAbs showed positive RBD-binding (**Extended Data Fig. 1c**). We then screened mAb clones showing high RBD ELISA positive rate for their neutralizing ability using an HIV- 1-based SARS-CoV-2 pseudovirus system and a VSV-based SARS-CoV-2 pseudovirus system (**Extended Data Fig. 2a-c**). Two mAbs, Clones 2 and 6, showed strongest binding and neutralizing activity (**Extended Data Fig. 2a-c**). As SARS-CoV-2 continues to mutate and evolve, leading to variants, it is critical to prevent viral escape from antibody recognition. To overcome this problem, antibody combinations from two or more mAbs have been developed and utilized ^42^. As an alternative, a single bispecific mAb can be used, as it can recognize two epitopes. One advantage of bispecific mAbs is that a single antibody product can be manufactured instead of two separate mAbs, which in theory could reduce the cost and formulation complexity. Thus, we generated a bispecific antibody using the antigen-specific variable regions of both Clones 2 and 6 (named as Clone 16). To generate the bispecific antibody, we utilized (i) a “knobs into holes” (KiH) methodology, and (ii) a CrossMAb-KiH bispecific backbone, to ensure the correct heterodimerization of the two different heavy chains, and (iii) a CH1 and CL exchange in heavy and light chain of one-half IgG to ensure correct pairing of heavy and light chains of each variable region (**Extended Data Fig. 1d**). We validated the antigen-specificity of the bispecific antibody along with its two parent mAbs using ELISA (**Extended Data Fig. 1e**).

To characterize the biophysical nature of the RBD reactivity of lead clones, we performed biolayer interferometry (BLI) and surface plasmon resonance (SPR) (**Fig. 1c-e**). BLI results showed that Clones 2 and 6 bound to the RBD with picomolar level dissociation constant (Kd) (**Fig. 1f**). The binding is particularly strong so that the dissociation stage was never observed in the BLI assay (**Fig. 1c**). SPR using an NTA-CHIP based assay with the RBD-antigen immobilized also revealed picomolar level dissociation constant (Kd) for Clones 2 and 6 (**Fig. 1d)**. In the NTA-CHIP, where the RBD-antigen is immobilized, multiple binding events can occur, leading to potential trimer formation of RBD protein and measurement of avidity. In parallel, we performed SPR with an Fc-CHIP based assay with pre-coated antibodies, which measures monovalent binding between RBD and the antibody. Results from Fc-CHIP SPR assay revealed strong affinity with single-digit nanomolar Kd of Clone 2 and Clone 6 (**Fig. 1e**). A humanized Clone 2 (named Clone 13) maintained strong binding affinity to RBD at low-double-digit nanomolar Kd (**Fig. 1e**). The bispecific Clone 16 also displayed high affinity to RBD also at single-digit nanomolar Kd (**Fig. 1e**), although not significantly higher than either parental clone alone.

### Cryo-EM structures of lead antibody clones bound to the SARS-CoV-2 spike define epitopes and binding conformations

We then generated Fab fragments of the lead clones and determined the cryo-EM structures of Clone 2 or 6 Fabs in complex with the ectodomain of SARS-CoV-2 spike trimer (S trimer) at ∼3 Å resolution (**Table 1**). In both cases, a S trimer is bound with three Fab molecules, one per RBD in various conformations. In total, we determined five different Fab-S trimer complex structures, each with the S trimer in a specific conformational state. Clone 2 Fab binds to S trimer in two states, one with 2 RBDs in the up conformation (60% of all complexes) and one with all 3 RBDs up (40%) (**Fig. 2a; Extended Data Fig. 3a**). Clone 6 Fab binds to S trimer in three states, one with all 3 RBDs down (26% of all complexes), one with 1 RBD up (43%) and one with 2 RBDs up (31%) (**Fig. 2b; Extended Data Fig. 3b**). Thus, Clones 2 and 6 Fab molecules are capable of recognizing SARS-CoV-2 S trimer in all possible RBD conformations, potentially reducing viral immune evasion when used in combination or as a bi-specific. Within each clone, the same Fab-RBD binding interface is maintained regardless of the RBD conformations across all S trimer states. Between the two clones, the Fab-RBD interfaces are different, although the two Fab clones bind RBD in similar locations with RBD adopting virtually the same conformation (RMSD 0.46 Å) (**Fig. 3a,b**). Clone 2 Fab-S trimer complexes appear more flexible than Clone 6 complexes, as indicated by less well-defined cryo-EM density in the Clone 2 Fab-RBD regions.

**Figure 2.**
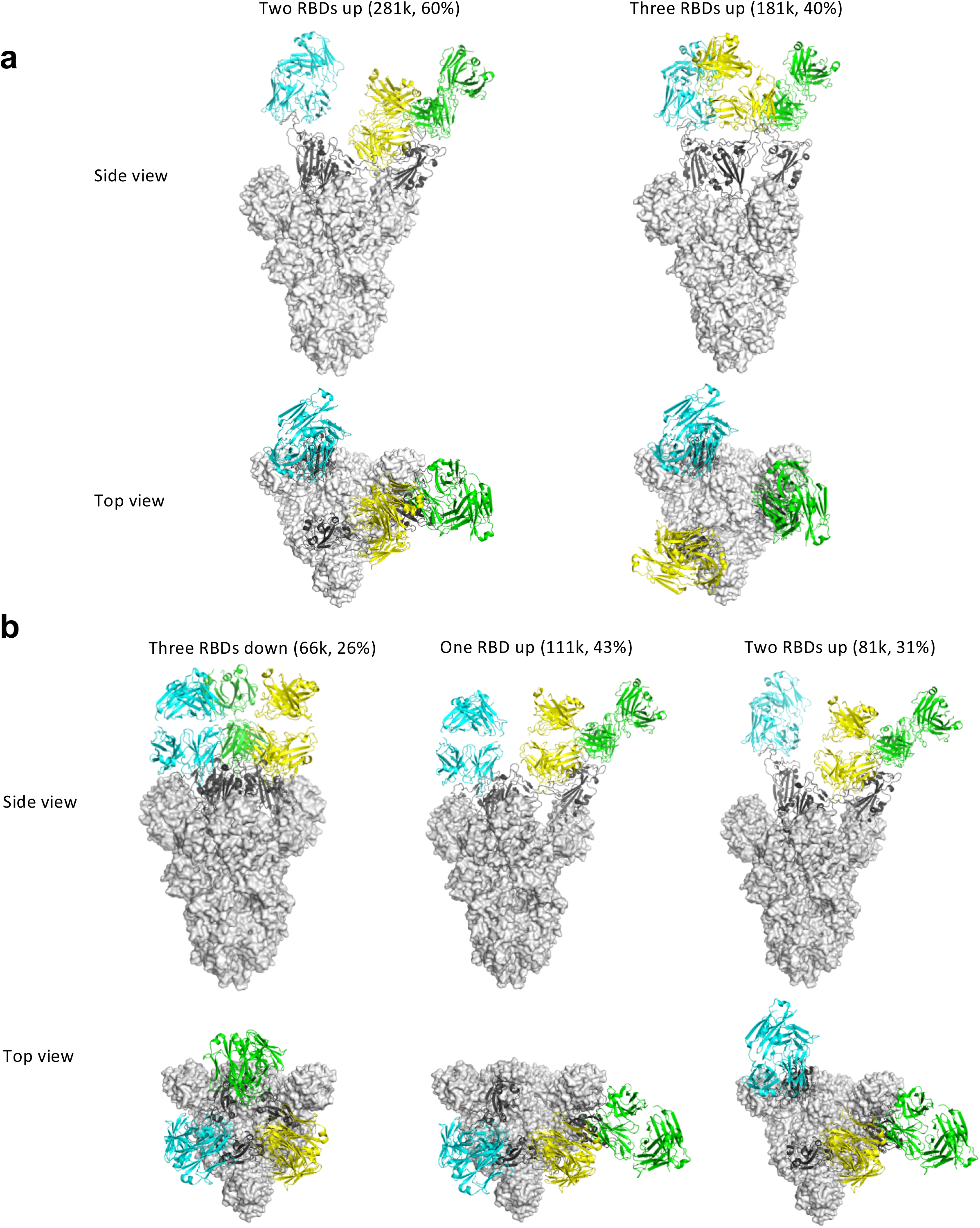
Cryo-EM structures of the ectodomain of SARS-CoV-2 spike trimer in complex with Clone 2 or Clone 6 Fab and Structural comparisons. **a-b**, Cryo-EM structures of the ectodomain of SARS-CoV-2 spike trimer in complex with Clone 2 **(a)** or Clone 6 **(b)**. Each Fab molecule is shown as ribbons in different colors, spike RBD is shown as dark gray ribbons, with the rest of spike trimer shown as gray surface. The particle distribution of each S trimer conformation is shown correspondingly.

**Figure 3.**
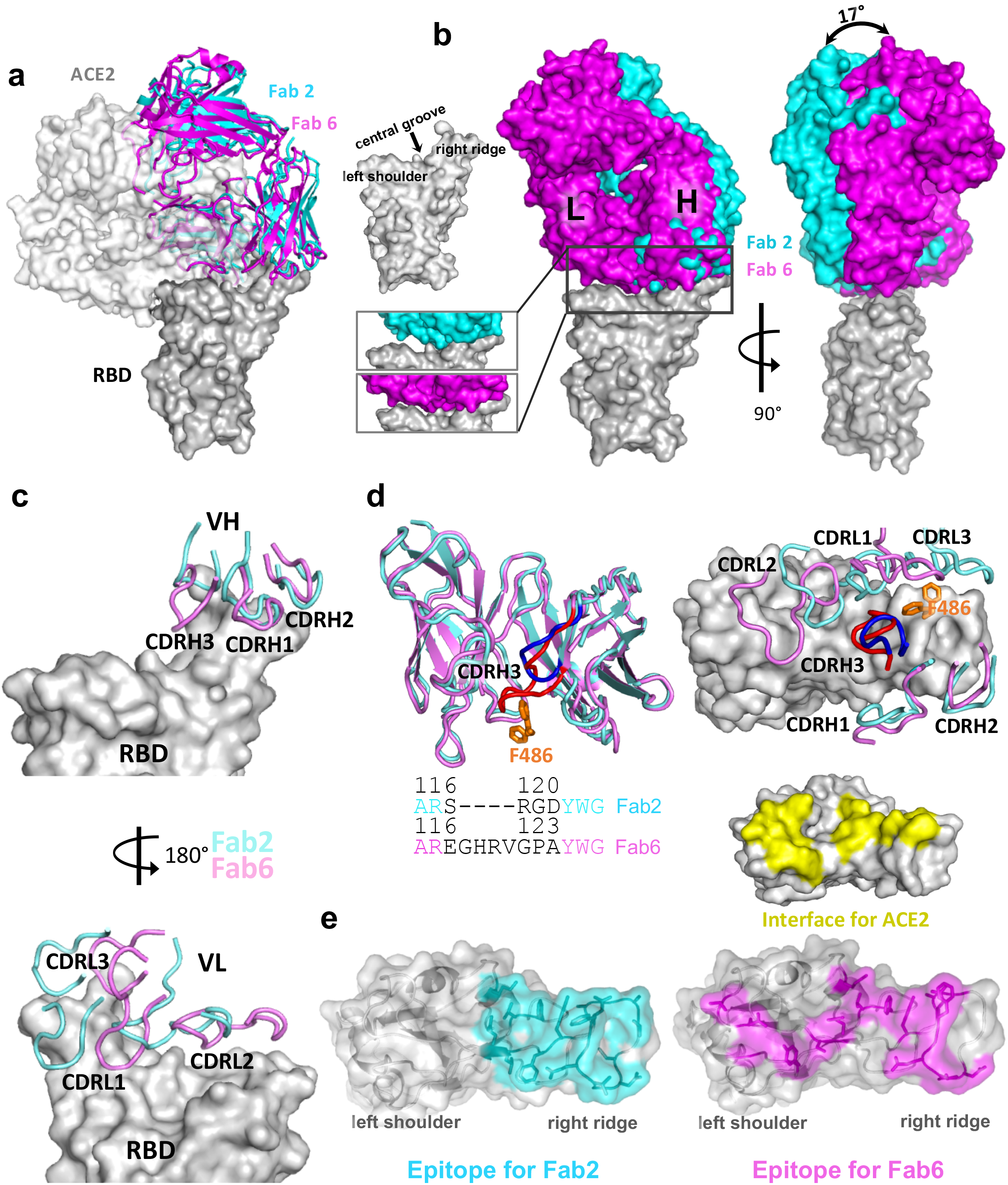
Binding interfaces and epitopes of lead monoclonal antibodies. **a**, Overlay of the structures of spike RBD from its complex with the host ACE2 receptor, Clone 2 or Clone 6 Fab indicates both antibody clones directly block the ACE2 access. **b**, Overlay of the structures of spike RBD from its complex with Clone 2 or Clone 6 Fab reveals a 17° rotation pivoted at the right ridge of spike RBD, resulting in less contact of Clone 2 Fab light chain with the left shoulder of spike RBD. The comparison of the Clone 2 and Clone 6 Fab binding interfaces with spike RBD is shown in the bottom parallel insets. The nomenclature of the top surface of spike RBD is described in the top inset. **c**, The orientations of the three CDRH (upper panel) and CDRL (lower panel) loops of Clone 2 or Clone 6 Fabs on spike RBD. The structures were overlaid with the RBD potions. **d**, Left, overlay of Clone 2 and Clone 6 Fab molecules alone. The CDRH3 loops that have the largest conformational difference are highlighted in brighter colors (blue for clone 2 and red for clone 6). Lower inset, amino acid sequence alignment of the CDRH3 loops of both clones. Right, spike RBD residue F486, which is shown as orange sticks, has a large conformational change upon RBD binding to the two clones. **e**, Comparison of spike RBD epitope regions for Clone 2 and Clone 6 Fabs, highlighted in cyan and magenta, respectively. The spike binding interface for hACE2 receptor is highlighted in yellow.

**Table 1.**
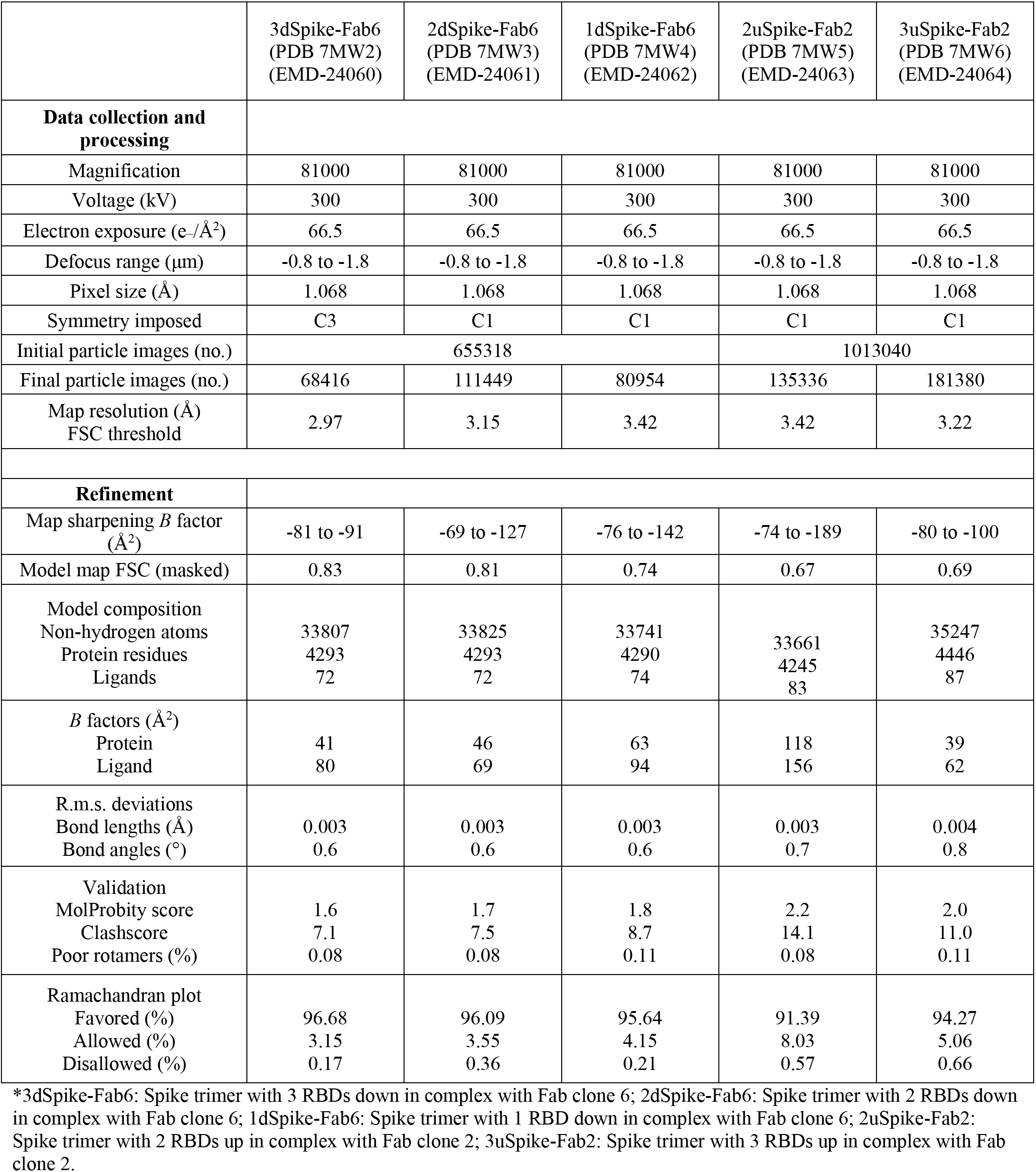
Cryo-EM Data Collection and Refinement Statistic.

Besides the major Fab-RBD interfaces described above, in the Clone 6 Fab complexes with 2 or 3 RBDs down, a down-RBD-binding Fab makes an additional contact to a side surface of an adjacent down-RBD to lock both RBDs in the closed conformation incompetent in ACE2 recognition (**Extended Data Fig. 4d left two panels**), which may provide a second neutralization mechanism. The additional contact is achieved by the CDRL1 loop of Clone 6 Fab that protrudes in between the two neighboring down-RBDs (**Extended Data Fig. 4c**), along with the framework region 3 (FWR3) of the Fab that interacts a nearby side surface of the adjacent down-RBD (**Extended Data Fig. 4d, left two panels**). In addition, in all complexes with 2 up/1 down-RBDs and the Clone 6 Fab complex with 2 down/1 up-RBDs, a down-RBD-binding Fab contacts the side surface of an adjacent up-RBD through another conserved interface mainly involving the CDRL1 loop of the Fabs (**Extended Data Fig. 4b; Extended Data Fig. 4d, right panels**), further stabilizing the conformation of the adjacent up-RBDs. These types of bivalent interactions are not observed for the up-RBD-binding Fabs.

Clone 2 and Clone 6 Fabs interact with spike RBD at similar locations, as both directly block access of the host receptor ACE2 (**Fig. 3a**). Clone 6 Fab shields the entire top surface of spike RBD (both “left shoulders” and “right ridge”), whereas Clone 2 Fab has a relative rotation of ∼17° pivoted on the RBD right ridge, resulting in a larger contact area there and less contact with the RBD left shoulder (**Fig. 3b**). When overlaying a Clone 2 Fab-bound down-RBD onto a Clone 6 Fab-bound down-RBD in the S trimer with more than one RBDs down, spatial clashes are detected between the Clone 2 Fab and the neighboring down RBD (**Extended Data Fig. 4a**). This also explains why we did not observe Clone 2 Fab-bound S trimer in a conformation state with more than one RBD down. In each case, all six complementarity determining regions (CDRs) of the Fab participate in RBD interactions (**Fig. 3c**). The 3 CDRH loops of the two clones share similar overall conformations and interact mainly with the right ridge of spike RBD (**Fig. 3c, upper panel**). The 3 CDRH loops of Clone 2 Fab form a large positively charged paratope, complementing a negatively charged epitope contributed by the right ridge of spike RBD (**Extended Data Fig. 5a**, upper panels). The 3 CDRH loops of Clone 6 Fab engage spike RBD predominantly through hydrophobic interactions (**Extended Data Fig. 5b**, upper left), with the CDRH3 loop also engaging RBD through modest electrostatic interactions (**Extended Data Fig. 5b,** upper right).

In contrast to the relatively conserved binding locations by the CDRH loops, the 3 CDRL loops of Clones 2 and 6 diverge in RBD-binding paratopes (**Fig. 3c, lower panel**). CDRL1 and CDRL3 of Clone 2 Fab primarily contact the epitopes at RBD right ridge and the CDRL2 loop engages RBD central groove, whereas CDRL3, CDRL1 and CDRL2 of Clone 6 Fab complement with the RBD right ridge, central groove and left shoulder, respectively, together covering the top surface of the RBD. Both Clones 2 and 6 Fab CDRL loops contact spike RBD primarily through hydrophobic interactions along with modest electrostatic interactions (**Extended Data Fig. 4b and 5a**, lower panels). A comparison of the Clone 2 and Clone 6 Fabs revealed that the most different region comes from the CDRH3 loop (**Fig. 3d**). The Clone 6 Fab contains a longer CDRH3 loop, which inserts into the central groove of spike RBD to generate a tighter engagement. As a consequence, this induces a substantial change in the RBD, which otherwise maintained a nearly identical conformation, where residue F486 flips to contact the CDRL3 loop of Clone 6 instead of interacting with the CDRH3 loop of Clone 2 (**Fig. 3d**). Even though the two Fabs interact with the RBD with similar buried surface areas (949 A2 in Clone 2 vs.894 A2 in Clone 6), the further spread-out RBD-binding mode of Clone 6 Fab is more similar to that of host ACE2 receptor (**Fig. 3e**).

Multiple mAbs have been generated to date and are in various stages of research and development, with some granted EUA for clinical use^3, 28, 43–45^. We compared our structures to several of the previously published mAb:Spike structures, and found that the RBD-binding modes of Clone 2 and Clone 6 Fabs differ from these SARS-CoV-2 neutralizing antibodies reported in primary literature and the protein databank (PDB). Three previously reported antibodies (2H2, CV05-163, and S2H13) target the spike RBD in somewhat similar orientations but with substantial rotations or shifts 46-48 (**Extended Data Fig. 6a**).Another two reported antibodies (CT-P59 and BD23) adopt RBD-binding conformations resembling those of Clone 2 and 6 Fabs ^49, 50^; however, the binding positions of heavy chains and light chains are exchanged (**Extended Data Fig. 6b**).

### Lead antibody clones potently neutralize SARS-CoV-2 WA1 and B.1.617 variant

We next tested the neutralization ability of the monospecific and bispecific clones. Because spike protein can mediate receptor-mediated membrane fusion^51^, we utilized a cell fusion assay in which expression of Wuhan- 1/WA1 SARS-CoV-2 spike on one cell can induce fusion with hACE2-expressing cells. All three antibodies (Clone 2, Clone 6 and Clone 16) effectively inhibited this spike mediated cell fusion activity (**Extended Data Fig. 2d-e**). Next, we used HIV-1-based pseudovirus system pseudotyped with SARS-CoV-2 Wuhan-1/WA1, B.1.351 and B.1.617 to evaluate neutralization potential of these mAbs. All three antibodies (Clone 2, Clone 6 and Clone 16) potently inhibited SARS-CoV-2-Wuhan-1 pseudovirus with low ng/mL level IC50 values (**Fig. 4a**). Due to the partial overlap in the binding domains in RBD, the bi-specific clone 16 was not stronger than either Clone 2 or Clone 6. These three antibodies also inhibited B.1.351 SARS-CoV-2 pseudovirus, with somewhat reduced potency (higher level of IC50 values, mid ng/mL for Clones 2 / 6, and high ng/mL for Clone 16) (**Fig. 4b**). In comparison, all three antibodies maintained potent neutralizing activity against the B.1.617 pseudovirus, with sub-single-digit to low-single-digit ng/mL level IC50 values (**Fig. 4c**). We also performed these experiments with Clones 2 and 6 as a cocktail combination (**Fig. 4d-f**). Although the combination is not superior to the single clone alone, the results again confirmed the potency of both Clones 2 and 6 as the single agents, where both clones showed strong potency against Wuhan-1/WA1 spike and B.1.617 pseudoviruses (single-digit ng/mL IC50) and slightly reduced potency against B.1.351 (**Fig. 4d-f**).

**Figure 4.**
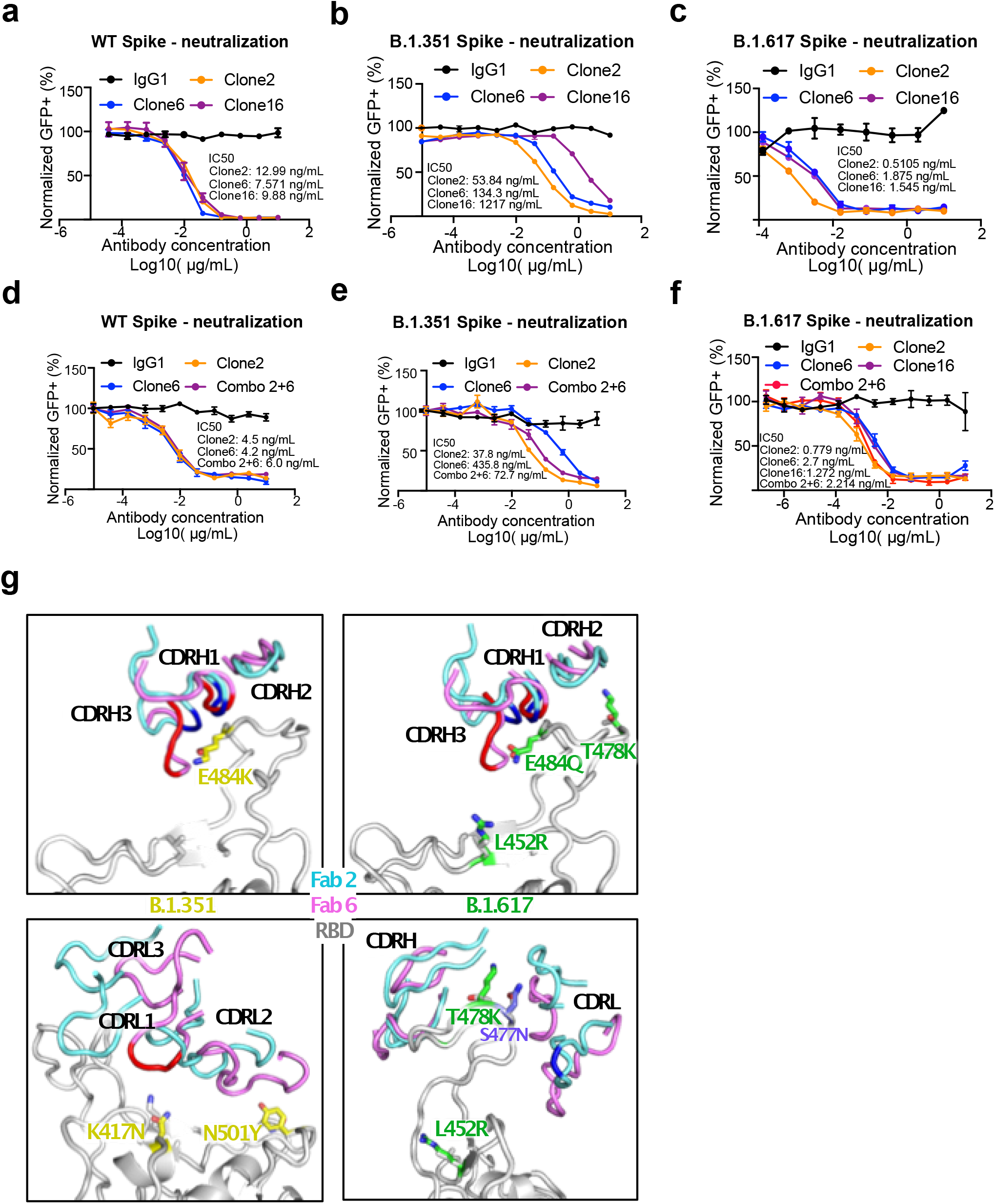
Lead mAb clones showed strong neutralization potency against WT/WA1 and B.1.617, with epitope and mutation analysis of the Fab-spike RBD interfaces. **a**, Neutralization assay on top mAb clones using WT/WA1 SARS-CoV-2 Spike pseudotyped HIV1-lentivirus carrying an EGFP reporter (pseudovirus). **b**, Neutralization assay on top mAb clones using B.1.351 variant SARS-CoV-2 Spike pseudovirus. **c**, Neutralization assay on top mAb clones using B.1.617 variant SARS-CoV-2 Spike pseudovirus. **d**, Neutralization assay on top mAb clones and their combination using WT/WA1 SARS-CoV-2 Spike pseudovirus. **e**, Neutralization assay on top mAb clones and their combination using B.1.351 variant SARS-CoV-2 Spike pseudovirus. **f**, Neutralization assay on top mAb clones and their combination using B.1.617 variant SARS-CoV-2 Spike pseudovirus. **g**, Structural analysis of the spike RBD mutations from the B.1.351 (left panels) and B.1.617 (right panels) variants at the interfaces with Clone 2 and Clone 6 Fabs. The residues mutated in spike RBD are shown as gray sticks in wild-type (WT) spike and yellow in the B.1.351 variant and green in the B.1.617 variant. Another frequent RBD mutation S477N in SARS-CoV-2 strains is also shown as slate sticks. The spike RBD is shown as light gray ribbon, and the CDR loops of Clone 2 and Clone 6 Fabs are shown as cyan and magenta ribbons, respectively, with the WT RBD-interacting residues highlighted in brighter blue and red colors

### Epitope and mutation analysis of the mAb:RBD structures on binding interface between the lead mAbs and hotspot mutations in Beta and Delta variants

To evaluate the structural impacts of the RBD mutations on binding of the Beta and Delta variants, we used our Spike:mAb Cryo-EM structures and generated homology models of the two variants (**Fig. 4g**). The Beta variant encodes three mutations (K417N, E484K, and N501Y) in spike RBD. Although N501 is not located within the epitope region, the Y501 mutation may have weak interactions with the CDRL2 loop of both Fab clones (**Fig. 4g**, lower left). K417 has no contact with Clone 2 Fab, but interacts with the CDRL1 loop of Clone 6 Fab, and the K417N mutation would likely abolish this interaction (**Fig. 4g**, lower left). E484, which is located on top of the RBD right ridge, is targeted by the CDRH1 and CDRH3 loops of both Fab clones, and has a larger contact area on Clone 6 Fab (**Fig. 4g**, upper left). The Delta variant encodes an E484Q at the position, which likely results in less disruption of its interaction with the Fabs (**Fig. 4g**, upper right). The two other mutation sites, L452 and T478, are not located within the antibody-binding surface (**Fig. 4g**, right). However, the T478K mutation is located close to the interface, especially that with Clone 2 Fab, and could affect RBD recognition by both clones (**Fig. 4g**, lower right).

### *In vivo* prophylactic and therapeutic efficacy of the lead mAbs against authentic SARS-CoV-2 virus

We next evaluated the potency of the lead antibodies against authentic SARS-CoV-2 (WA1/2020) infection. All three antibodies (Clone 2, Clone 6 and the bispecific Clone 16) inhibited infection of SARS-CoV-2 WA1/2020 (low-mid ng/mL level IC50s) (**Fig. 5a**). We then assessed efficacy of mAbs against SARS-CoV-2 *in vivo*, when administered as either pre-exposure prophylaxis or post-exposure therapy (**Fig. 5b**). We performed protection studies with SARS-CoV-2 using K18-hACE2 transgenic mice ^52–54^. We challenged K18- hACE2 mice with 2 x 10^3^ plaque forming units (PFU) of SARS-CoV-2 WA1/2020 virus (**Fig. 5b**). The hACE2 transgenic mice were randomly divided into three group, and mice in each group were injected with 20 mg/kg of Clone 2 mAb, Clone 6 mAb, or placebo control; with treatment given as a single dose either 24 h before or 18 h after viral infection (**Fig. 5b**).

**Figure 5.**
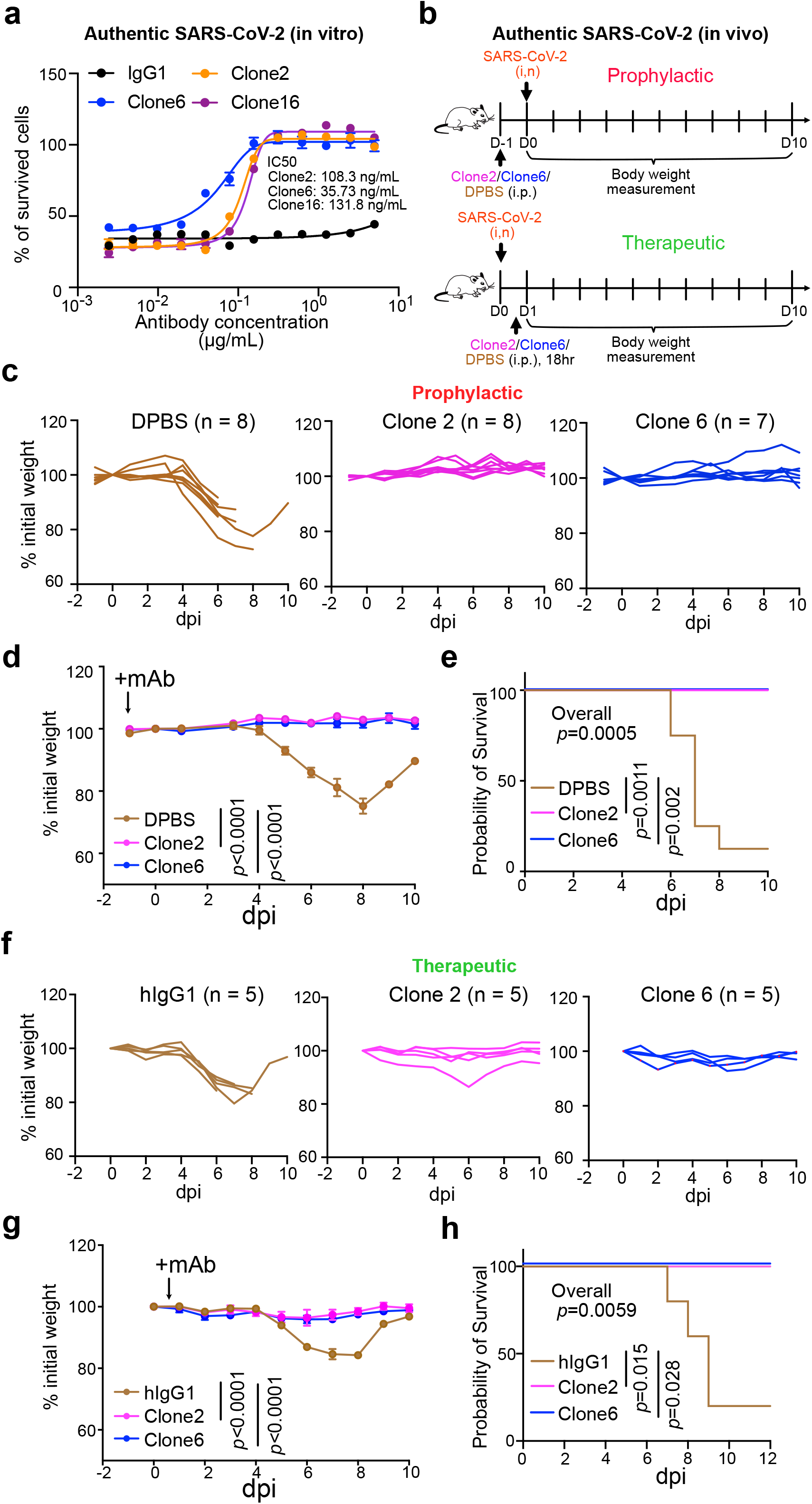
Lead monoclonal antibodies have potent prophylactic and therapeutic *in vivo* efficacy against replication competent SARS-CoV-2 virus. **a**, In vitro neutralization of top mAb clones against authentic SARS-CoV-2 in BL3 setting **b**, Schematics of *in vivo* efficacy testing of top mAb clones against lethal challenges of authentic SARS-CoV- 2 in hACE2 transgenic mice at BL3 level, in both prophylactic (upper panel) and therapeutic (bottom panel) settings. **c-e**, Prophylactic efficacy testing. **c and d**, Body weight curves of antibody and placebo (DPBS) treated hACE2 transgenic mice under lethal challenges of authentic SARS-CoV-2. **c**, Spider curves of body weight changes of individual mouse plotted separated by group. **d**, Mean +/- SEM curves of body weight changes. (Clone 2, n = 8; Clone 6, n = 7, and DPBS, n = 8). **e**, Survival curves of antibody and placebo (DPBS) treated hACE2 transgenic mice under lethal challenges of authentic SARS-CoV-2 (the same experiment in **c-d**). Therapeutic efficacy testing. **f-h**, Body weight curves of antibody and placebo (isotype hIgG1) treated hACE2 transgenic mice under lethal challenges of authentic SARS-CoV-2. **f**, Spider curves of body weight changes of individual mouse plotted separated by group. **g**, Mean +/- SEM curves of body weight changes. (Clone 2, n = 5; Clone 6, n = 5, and hIgG1, n = 5). **h**, Survival curves of antibody and placebo (hIgG1) treated hACE2 transgenic mice under lethal challenges of authentic SARS-CoV-2 (the same experiment in **f-g**). In this figure: Data are shown as mean ± s.e.m. plus individual data points in dot plots. Statistics: Two-way ANOVA was used to assess statistical significance for multi-group curve comparisons; Log-Rank test was used to assess statistical significance for survival curve comparisons; unless otherwise noted. The p-values are indicated in the plots. Source data and additional statistics for experiments are provided in a supplemental excel file.

In the prophylactic setting, all (8/8, 100%) mice in the placebo group developed severe disease due to viral challenge, and most (7/8, 87.5%) of them lost substantial body weight and succumbed from the disease, with only one mouse recovering weight (**Fig. 5c-e**). In contrast, all mice receiving treatment of either Clone 2 (8/8, 100%) or Clone 6 (7/7, 100%) maintained their body weight throughout the duration of the study and survived SARS-CoV-2 infection (**Fig. 5c-e**). In the therapeutic setting, all (5/5, 100%) mice in the placebo group developed severe disease, and most (4/5, 80%) of them lost body weight (**Fig. 5f-h**). In contrast, all mice treated with Clone 2 (5/5, 100%) or Clone 6 (5/5, 100%) at + 18 h maintained their body weight and survived the viral infection with no signs of disease (**Fig. 5f-h**). These data suggested that these mAbs can protect the animals from lethal SARS-CoV-2 infection in either prophylactic or therapeutic settings.

### Generation, biophysical characterization, and functional testing of a humanized mAb clone

To improve clinical translatability, we also developed a humanized Clone 2, using standard antibody humanization approaches with framework humanization and engineered mutations, based on canonical human antibody backbone sequences as well as the antibody : RBD CryoEM structures (Methods). A resultant humanized clone that maintains RBD specificity was generated (Clone 13A) (**Fig. 6a**). We purified Clone 13A with other lead clones for characterization and functional studies (**Fig. 6b**). BLI data showed that Clone 13A has a low double-digit Kd value (**Fig. 6c**).

**Figure 6.**
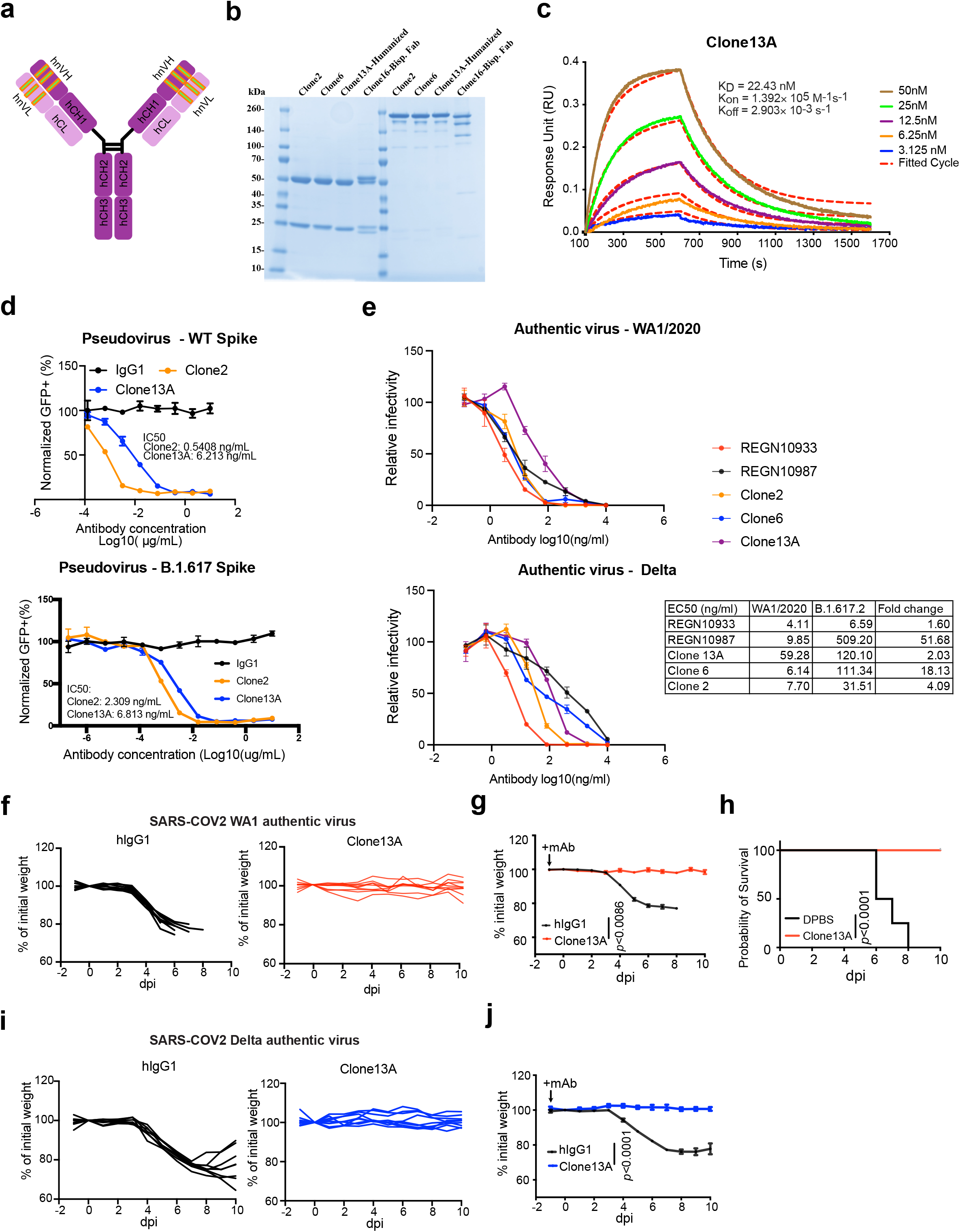
Generation, biophysical characterization, and functional testing of a humanized mAb clone. **a**, Schematics of antibody humanization. **b**, SDS-PAGE of purified antibodies of humanized clone (Clone 13A) along with other clones. **c**, Octet measurement of Clone 13A using the BLI assay. **d**, (Top) Neutralization assay on Clone 2 and a Clone 13A (humanized Clone 2) using WT SARS-CoV-2 Spike pseudotyped HIV-lentivirus carrying an EGFP reporter. (Bottom) Neutralization assay on Clone 2 and a Clone 13A (humanized Clone 2) using B.1.617 variant SARS-CoV-2 Spike pseudotyped HIV-lentivirus carrying an EGFP reporter. **d**, *In vitro* neutralization of top mAb clones (Clones 2, 6, 13A) along with representative therapeutic antibody clones (RGEN 10933, 10987) against authentic SARS-CoV-2 WA1/2020 and B.1.617.2 (Delta) viruses in BL3 setting. **f-g**, Body weight curves of humanized antibody Clone 13A and placebo (isotype hIgG1) treated hACE2 transgenic mice under lethal challenges of authentic SARS-CoV-2 WA1. **f**, Spider curves of body weight changes of individual mouse plotted separated by group. **g**, Mean +/- SEM curves of body weight changes. (Clone 13A, n = 9, and hIgG1, n = 8). **h**, Survival curves of Clone 13A antibody and placebo (hIgG1) treated hACE2 transgenic mice under lethal challenges of authentic SARS-CoV-2 (the same experiment in **f-g**). **i-j**, Body weight curves of humanized antibody Clone 13A and placebo (isotype hIgG1) treated hACE2 transgenic mice under lethal challenges of authentic SARS-CoV-2 Delta. **i**, Spider curves of body weight changes of individual mouse plotted separated by group. **j**, Mean +/- SEM curves of body weight changes. (Clone 13A, n = 8, and hIgG1, n = 8). In this figure: Data are shown as mean ± s.e.m. plus individual data points in dot plots. Statistics: Two-way ANOVA was used to assess statistical significance for multi-group curve comparisons; Log-Rank test was used to assess statistical significance for survival curve comparisons; unless otherwise noted. The p-values are indicated in the plots. Source data and additional statistics for experiments are provided in a supplemental excel file.

Clone 13A also potently neutralized WT/WA1 and B.1.617 pseudoviruses (single-digit ng/mL IC50) (**Fig. 6d**). We also performed authentic virus neutralization assays with Clone 13A, along with Clone 2, Clone 6 and two EUA antibodies (RGEN 10933 and 10987). Clones 2, 6 and 13A all potently neutralized SARS-CoV-2 WA1/2020 and Delta variant authentic viruses (**Fig. 6e**). Against Delta variant, RGEN 10933 has a 1.6x drop in potency, Clone 13A has a 2x drop, Clone 2 has a 4x drop, Clone 6 has an 18x drop, while RGEN 10987 has a 52x drop.

We also performed *in vivo* challenge experiment with Clone 13A using authentic viruses (**Fig. 6f-h**). Similar to the prior results all (8/8, 100%) mice in the placebo group developed severe disease due and succumbed from infection (**Fig. 6f,h**). In contrast, all mice receiving Clone 13A (9/9, 100%) maintained body weight throughout the duration of the study and survived from SARS-CoV-2 WA1 infection (**Fig. 6g,h**). We also tested Clone 13A against the Delta variant authentic virus challenge. All (8/8, 100%) mice in the placebo group developed severe disease due and succumbed from infection (**Fig. 6i**). In contrast, all mice receiving Clone 13A (8/8, 100%) maintained body weight throughout the duration of the study (**Fig. 6j**). Of note, the Delta virus appeared to be not lethal for hACE2 mice within the experimental setting, thus the survival benefit can not be measured against Delta. These data confirmed that the humanized Clone 13A maintained its potency and protective activity *in vivo* against authentic SARS-CoV-2, both the original WA1 and Delta variant in the B.1.617 lineage.

## Discussion

The ongoing COVID-19 pandemic and continued emergence of SARS-CoV-2 variants has necessitated the rapid development of therapeutic interventions. The discovery and development of neutralizing antibodies with expanding collections of epitopes are critical to provide countermeasure options against escape seen with emerging variants of concern. We combined SARS-CoV-2 Spike RBD protein immunization with high-throughput single cell BCR sequencing technology to establish a platform to develop neutralizing antibody candidates. We identified two highly potent and specific SARS-CoV-2 neutralizing mAb clones with single-digit nanomolar affinity and low-picomolar avidity. We also generated a bispecific antibody of these two lead clones, as well as a potent humanized clone. The lead antibodies showed strong neutralization ability against SARS-CoV-2 and the highly transmissible B.1.617 lineage that poses a risk of reducing the efficacy of currently available therapeutic antibodies and prophylactic vaccines.

The SARS-CoV-2 spike protein is dynamic, and three conformations of spike prefusion trimers have been detected on intact virions: all RBDs down, 1 RBD up, and 2 RBDs up, with the last one only existing in vitro with multiple stabilizing mutations ^55^. The spike RBD needs to extend upwards to be accessible to ACE2, and the presence of host ACE2 changes the population distribution of different spike conformations by promoting the RBD towards the up/open state favorable for ACE2 binding ^56^. Genetic variations in spike can also change the conformational equilibrium of the trimer, subsequently affecting virus infectivity57. Although antibody therapy has been developing rapidly prevent or treat SARS-CoV-2, infection, the precise determinants of neutralization potency remain unclear. In addition to direct receptor blockade by antibody binding, modulation of spike-mediated membrane fusion by altering the ACE2-triggered spike protein conformational cycle has been suggested as another determinant of the antibody neutralization potency^51^.

Depending on the binding mode, some antibodies may facilitate the spike conformational cycle to the final stage with all 3 RBDs open, or lock the spike trimer in a pre-fusion state, therefore enhancing or inhibiting the cell membrane fusion and syncytium formation. Based on our cryo-EM analysis, Clone 6 Fab binds to spike trimer preferentially with at least one RBD down, and effectively skews the spike trimer towards pre-fusion states (**Fig. 2b**). Besides making two neighboring down-RBDs inaccessible for ACE2 binding (**Extended Data Fig. 4c and 4d**, left panels), a down-RBD-binding Clone 6 Fab can also interact with an adjacent up-RBD through a quaternary epitope located at the side wall of the up-RBD (**Extended Data Fig. 4d**, right), which clashes with ACE2 binding (**Extended Data Fig. 4e**, right), thereby directly blocking ACE2 access to two RBDs simultaneously. This bipartite binding mode presumably is more stable than the single binding mode with the up-RBD alone, explaining why no spike trimer with all 3 RBDs open was detected when complexed with Clone 6 Fab. We hypothesize that the Clone 6 Fab binding to this secondary epitope of RBD helps lock the spike trimer in the pre-fusion form, which inhibits the spike-mediated cell membrane fusion by historical virus and even the B.1.1.7 variant that has enhanced binding affinity to ACE2 58 (**Extended Data Fig. 2d-e**,middle). In contrast, due to the different binding conformations of Clone 2 Fab on spike RBD, the spike trimer has been detected in skewed states that favorite RBDs up (**Fig. 2a**), which mimics the effect of ACE2 binding during the conformational cycle of the spike trimer. Nonetheless, we found that Clone 2 Fab also efficiently determining the neutralization potency of spike-targeting antibody is complex.

Antibody resistance of SARS-CoV-2 B.1.1.7, B.1.351, and B.1.617 lineage variants has been reported ^7, 14, 59^. The B.1.1.7 variant of SARS-CoV-2 spike contains a single mutation (N501Y) in the RBD, which has been reported to enhance the spike RBD-ACE2 binding affinity that could disfavor antibody neutralizations competing with ACE2 for RBD binding ^58^. N501 does not directly interact with either Clone 2 or Clone 6 Fab, although the mutation may generate allosteric effects on other CDRL loops nearby to disrupt the binding interface ^58^. While the B.1.1.7 variant spreads faster, has higher case-fatality rate, and some antibody resistance, it does not reduce the efficacy of the currently approved vaccines ^7, 60–62^. In contrast, the B.1.1351 variant of SARS-CoV-2 showed increased rate of transmission, resistance to antibody therapeutics, and reduced vaccine efficacy ^6–8, 61^. In addition to the N501Y mutation, the B.1.351 variant has two additional point mutations K417N and E484K in spike RBD, which perturbs the RBD epitope recognition by both antibody clones, explaining their reduced potencies against the B.1.351 variant. The lineage B.1.617 has mutations in spike including G142D, E154K, L452R, E484Q, D614G, P681R, and Q1071H 15, which could affect a number of leading therapeutic antibodies tested to date 14,63,64. The L452R variant evades cellular immunity and increases infectivity 65. L452R and S477N might affect the potency of Clones 2 and 6 to some degree based on the structure.

Several therapeutic antibodies have been granted EUA for clinical use by the FDA, such as two from Regeneron developed in a previous study^66^. Recent studies showed that some of the EUA mAbs have significant reduction in neutralization activities against B.1.617 lineage variants^14, 63^. Many antibodies have been developed by the field and tested against various VoCs^64, 67, 68^. Our data also showed that RGEN 19087 has substantially diminished potency against Delta. To better understand impact of Delta mutations on antibody neutralization potency, we performed structural; analysis with the Delta RBD (**Extended Data Fig. 8**). The epitopes of REGN-10987 centered around the RBD left shoulder loop on the β1 end, which hereafter we termed left shoulder loop 1 (**Extended Data Fig. 8**). Clone 2 primarily targets a region at the right ridge of RBD. The L452R mutation of Delta is on the C-terminus edge of the left shoulder loop 1 and is critical for the loop 1 conformation. L452R is observed to favor an orientation interacting with the Y351 and is likely to disrupt the original conformation of left should loop 1, consistent with the reduction of the affinity and potency of the REGN-10987 antibody. The E484Q, on the other hand, was not observed to significantly change side chain orientation and is less likely to cause drastic conformational changes on nearby loops. The mutation induced side-chain orientation as well as nearby loop conformation change plus distinct antibody targeting epitopes are consistent with the observation that Clone 2 retained its neutralization potency against Delta, whereas REGN-10987 did not.

In summary, in a concerted global effort against COVID-19, a number of candidate therapeutics, antibodies, and vaccines have been developed to date ^69^. Our lead antibody clones are distinct different from existing antibodies reported to date in their binding geometry and footprints, are highly inhibitory against SARS-CoV- 2 and several variants of concerns, particularly the Delta variant from the B.1.617 lineage. These antibodies thus expand the repertoire of COVID-19 countermeasures against the SARS-CoV-2 pathogen and its emerging and potentially more dangerous variants.

## Acknowledgments

We thank various members from Chen, Xiong and Wilen labs for discussions and support. We thank Dr. Bieniasz for providing pseudovirus reporter plasmids. We thank staffs from various Yale core facilities (Keck, YCGA, HPC, Biophysics, YARC, Cryo-EM, CBDS and others) for technical support. We thank various support from Departments of Genetics, MBB, Laboratory Medicine, Immunobiology, Internal Medicine and Pharmacology; Institutes of Systems Biology and Cancer Biology; Dean’s office of Yale School of Medicine and the office of Vice Provost for Research.

This work is supported by DoD PRMRP IIAR (W81XWH-21-1-0019) and discretionary funds to SC; discretionary funds to YX; Ludwig Foundation, Mathers Foundation, Burroughs Wellcome Fund, NIH K08 AI128043, NIH R01 AI148467 to CBW; NIH R01 AI157155 to MSD; NIH/NIAID R01 AI150334 to RS. YCGA / HPC were supported by NIH Award 1S10OD018521. The T200 Biacore instrumentation was supported by NIH Award S10RR026992-0110.

## Methods

### Institutional Approval

This study has received institutional regulatory approval. All recombinant DNA (rDNA) and biosafety work were performed under the guidelines of Yale Environment, Health and Safety (EHS) Committee with approved protocols (Chen-15-45, 18-45, 20-18, 20-26; Xiong-17302; Wilen-18/16-2). All animal work was performed under the guidelines of Yale University Institutional Animal Care and Use Committee (IACUC) with approved protocols (Chen-2018-20068; Chen-2020-20358; Wilen-2018-20198).

### Animal immunization

Standard 28-day repetitive immunization protocol was utilized to for immunization. *M. musculus* (mice), 6-12 weeks old females, of C57BL/6J and BALB/c strains, were used for immunization. First, all mice are ear- marked and around 200 μl blood was taken as pre-immunization sample, where serum was collected from the blood by centrifugation (1,000 *g* for 10 min). Two days later (day 0), for each mouse, 20 μg SARS-CoV-2 RBD-his tag protein (Sino biological) in 100 μl PBS was mixed with 100 μl Complete Freund’s Adjuvant (CFA) with 3-way stop Cock. The fully emulsified mixture was subcutaneously injected into the back of each mouse. On day 7, a second immunization was performed, where each mouse was injected subcutaneously with 20 μg RBD-his tag protein fully emulsified with Incomplete Freund’s Adjuvant (IFA). On day 13, around 50 μl of blood from each mouse was obtained for serum preparation as first bleeds. On day 14, a third immunization is performed, where all the procedures were similar to the second immunization. On day 20, second bleeds were taken. On day 21, forth immunization is performed, where all the procedures were similar to the second immunization. On day 24, each mouse receives 20 μg RBD-his tag protein in 200 μl PBS intraperitoneally as final immunization. On day 28, mice with strong serum conversion detected by ELISA were sacrificed. Spleen, lymph nodes, and bone marrow were collected for B cells isolation and purification for single cell BCR sequencing. Serums from pre, first and second bleeds were subjected to ELISA for anti- RBD tilter determination.

### Mouse B cell isolation and purification

Primary B cells from spleen, draining lymph nodes, bone marrow of RBD-his tag protein immunized mice were isolated and purified with mouse CD138 MicroBeads (Miltenyi Biotec, 130-098-257) following standard protocol provided by the manufacturer. Spleens and draining lymph nodes were homogenized gently. Bone marrows were fragmented, rinsed with PBS containing 2% FBS, and filtered with a 100 μm cell strainer (BD Falcon, Heidelberg, Germany). The cell suspension was centrifuged for 5 min with 400 *g* at 4 °C. Erythrocytes were lysed briefly using ACK lysis buffer (Lonza) with 1mL per spleen for 1∼2 mins before adding 10 mL PBS containing 2% FBS to restore iso-osmolarity. The single-cell suspensions were filtered through a 40 μm cell strainer (BD Falcon, Heidelberg, Germany). CD138 positive B cells were isolated using magnetic cell sorting by positive selection according to the manufacturer’s instructions. Cell samples post-magnetic selection were counted and prepared for single cell BCR sequencing.

### Single cell BCR sequencing

The enriched CD138^+^ plasma cells and progenitor B cells were loaded on a 10X Chromium Next GEM Chip G. The target cell number was 10,000 cells per sample. Single-cell lysis and RNA first-strand synthesis were performed using Chromium Next GEM Single Cell 5’ Gel Bead V3.1 according to the manufacturer’s protocol. The following RNA and V(D)J library preparation was performed according to the manufacturer’s protocol (Chromium Next GEM Single Cell V(D)J reagent kit, mouse BCR). The resulting VDJ-enriched libraries were sequenced following the reading mode recommended by 10x Genomics. Sequencing was performed on a NovaSeq targeted for 10,000 reads/cell, with a total of 100 million reads.

### Single cell VDJ sequencing data analysis

Raw sequencing data were processed using Cell Ranger v3.1.0 with default settings, aligning the reads to the GRCm38 mouse VDJ reference. Outputs from Cell Ranger were then visualized using the Loupe V(D)J Browser for quality control assessment and to identify the top enriched clonotypes. The consensus amino acid sequences for the top-ranked heavy/light chain pairs in each sample were then extracted and codon-optimized for human expression.

### Plasmid construction

The cDNA sequences of the paired variable heavy and light chain region of anti-RBD antibody clones were synthesized as gBlocks (IDT) and cloned by the Gibson assembly (NEB) into human IgG1 heavy chain and light chain expression plasmids, pFUSEss-CHIg-hG1(InvivoGen, pfusess-hchg1) and pFUSE2ss-CLIg-hK (InvivoGen, pfuse2ss-hclk), respectively. pFUSEss-CHIg-hG1 plasmid is a cloning plasmid that expresses the constant region of the human IgG1 heavy chain and includes multiple cloning sites to enable cloning of the heavy chain (CH) variable region. Parallelly, pFUSE2-CLIg-hK is a cloning plasmid that expresses the constant region of the human kappa light chain and contains multiple cloning sites to enable cloning of the light chain variable region. For anti-RBD antibody clones’ heavy chain plasmid cloning, gBlocks, containing cDNA sequence of variable region of heavy chain of anti-RBD antibody clones and the regions overlapping with corresponding flanking sequences of EcoRI and NheI restriction sites pFUSEss-CHIg-hG1, were ordered from IDT. pFUSEss-CHIg-hG1 were digested with EcoRI and NheI restriction enzyme (ThermoFisher). These synthesized gBlocks were cloned into gel-purified restriction enzyme digested backbone by the Gibson assembly (NEB). For anti-RBD antibody clones’ light chain plasmid cloning, gBlocks, containing cDNA sequence of variable region of light chain of anti-RBD antibody clones and the regions overlapping with corresponding flanking sequences of EcoRI and BsiWI restriction sites pFUSE2ss-CLIg-hK, were ordered from IDT. The gBlocks were then cloned into the pFUSE2ss-CLIg-hK backbone, which was digested with EcoRI and BsiWI restriction enzyme (ThermoFisher).

The bi-specific antibody with the same Fab regions of clone 2 and clone 6 was generated by using the CrossMab-KiH bi-specific constructs ^70^. The CrossMab-KiH bi-specific constructs were designed and generated basing on pFUSEss-CHIg-hG1 and pFUSE2ss-CLIg-hK. The bi-specific antibody consists of two hetero-half IgG1, one is knob IgG1 and the other is hole IgG1 (Knob-in-Hole conformation). Four plasmids were employed: pFUSE2ss-knobLight-hK, pFUSE2ss-knobheavy-hG1, pFUSE2ss-HoleLight-hK, pFUSE2ss-HoleHeavy-hG1. The pFUSE2ss-knobLight-hK is pFUSE2ss-CLIg-hK with no further editing. The pFUSE2ss-knobheavy-hG1 contains two knob mutations (T366W, S354C) in the CH3 region when compared with pFUSEss-CHIg-hG1. The gBlock (pPR024), containing constant region of heavy chain with two knob mutations and the regions overlapping with corresponding flanking sequences of NsiI and NheI restriction sites in pFUSEss-CHIg-hG1 was ordered from IDT, and then cloned into NsiI and NheI restriction enzymes digested pFUSEss-CHIg-hG1 backbone by the Gibson assembly (NEB). The pFUSE2ss-HoleLight- hK was generated by replacing the constant region of Light chain (CL) in pFUSE2ss-CLIg-hK with CH1 region of heavy chain in pFUSEss-CHIg-hG1 vector. The CH1 region were PCR amplified from pFUSEss-CHIg-hG1vectors with a forward primer (oPR81-F) and a reverse primer (oPR82-R) containing regions overlapping with corresponding flanking sequences of the and NcoI and NheI restriction sites in the pFUSE2ss-CLIg-hK. CH1 PCR amplified fragments were gel-purified and cloned into restriction enzyme digested pFUSE2ss-CLIg-hK by the Gibson assembly (NEB). The pFUSE2ss-HoleHeavy-hG1 possesses three “hole” mutations (T366S, L368A, Y407V) in the CH3 region and a Y349C on the “hole” side to form a stabilizing disulfide bridge. In addition, to get the correct association of the light chain and the cognate heavy chain, the CH1 region in the pFUSE2ss-HoleHeavy-hG1 was exchanged with constant region of light chain (CrossMab conformation). The gBlock (pPR023), containing cDNA sequence of constant region of light chain, CH2 and CH3 with “hole” mutations, and regions overlapping with corresponding flanking sequences of NsiI and NheI restriction sites in pFUSEss-CHIg-hG1 was ordered from IDT, and cloned into NsiI and NheI restriction enzymes digested pFUSEss-CHIg-hG1 backbone through Gibson assembly (NEB). All plasmids were sequenced and Maxiprepped for subsequent experiments.

### Cloning of SARS-CoV-2 spike variants

The construct of wild-type (WT) SARS-CoV-2 ectodomain of spike trimer is a gift from Dr. Jason S. McLellan at University of Texas at Austin ^34^. The recently emerged SARS-CoV-2 spike SA variant B.1.351 ^7^ and Indian variant B.1.617 ^11^ was generated by standard cloning. The pVP21-SA variant includes four mutations in the N-terminal domain (L18F, D80A and D215G, R246I), three mutations at key residues in the RBD (N501Y, E484K and K417N), and one is in loop 2 (A701V). The pVP28-Indian variant includes seven mutations in Spike G142D, E154K, L452R, E484Q, D614G, P681R, Q1071H.The pVP21-SA and pVP28-Indian were generated based on pcDNA3.1-pSARS-CoV-2-S, which was derived by insertion of a synthetic human codon-optimized cDNA (Geneart) encoding a WA1 SARS-CoV-2 S protein. For pVP21-SA-variant, two gBlocks, contains mutations in SA variant regions overlapping with corresponding flanking sequences of NheI and BsrGI restriction sites pcDNA3.1-pSARA-CoV-2. The gBlocks were then cloned into pcDNA3.1-pSARA- CoV-2 backbone, digested with NheI and BsrGI restriction enzyme (Thermo fisher) through Gibson assembly. For pVP28-Indian, four gBlocks, contains mutations in Indian variant regions overlapping with corresponding flanking sequences of NheI and BamHI restriction sites pcDNA3.1-pSARA-CoV-2. The gBlocks were then cloned into pcDNA3.1-pSARA-CoV-2 backbone, digested with NheI and BamHI restriction enzyme (Thermo fisher) through Gibson assembly. For the HIV1-based SARS-CoV-2 spike pseudotyped virus generation, WT pcDNA3.1-pSARS-CoV-2-S, pVP21-SA-variant, and pVP28-Indian variant lacking the C-terminal 19 codons were employed. A pair of forward and reverse primers were utilized to amplify fragments lacking the C-terminal 19 codons with pVP21-SAvariant and pVP28-Indian variant as template separately. The amplified fragments were gel-purified and cloned into pVP21-SAvariant backbone and pVP28-Indianvariant backbone, digested with BbvCI and BamHI.

### Cell culture

HEK293FT (ThermoFisher) and 293T-hACE2 (gifted from Dr. Bieniasz’ lab) cell lines were cultured in complete growth medium, Dulbecco’s modified Eagle’s medium (DMEM; Thermo fisher) supplemented with 10% Fetal bovine serum (FBS, Hyclone),1% penicillin-streptomycin (Gibco) (D10 media for short). Cells were typically passaged every 1-2 days at a split ratio of 1:2 or 1:4 when the confluency reached at 80%. Expi293F^TM^ (ThermoFisher) cells were cultured in Expi293™ Expression Medium (ThermoFisher) in 125- mL shaker flask in a 37 °C incubator with 8% CO2 on an orbital shaker rotating at 125 rpm. For routine maintenance, Expi293F^TM^ cells were grown to 3-5×10^6^ cells/mL, then split to 0.3-0.5×10^6^ cells/mL every 3 days.

### Expression and purification of WT SARS-CoV-2 ectodomain of spike trimer

The WT ectodomain of SARS-CoV-2 spike trimer was expressed in Expi293F cells. For 100 mL expression scale, 100 μg construct DNA was mixed with 400 μg polyethylenimine in 10 mL Opti-MEM® I Reduced-Serum Medium (ThermoFisher) for 30 mins, and then added into 90 mL Expi293F cells at a density of 2.5- 3×10^6^ cells/mL for incubation, shaking at 125 rpm in a 37 °C incubator with 8% CO2. After 5 days, the medium with the secreted protein was harvested and loaded onto an ion exchange column. Fractions containing the target protein was pooled and further purified using a Ni-NTA affinity column, followed by size exclusion chromatography using a Superose 6 10/300 column (GE Healthcare) with a buffer of 30 mM Tris, pH 8.0, 100 mM NaCl. The monodispersed peak containing the ectodomain of the spike trimer was pooled and concentrated for subsequent analysis.

### Recombinant antibody generation

The top-ranked enriched IgG clones were selected and cDNAs of relative variable region of paired heavy- and light-chain were codon-optimized and cloned separately into human IgG1 heavy chain and light chain expression vectors, containing the human IgG1 constant regions (pFuse plasmids). IgG1 antibodies were expressed in Expi293F^TM^ cells. ExpiFectamine 293 transfection kit (Thermo fisher) was utilized for heavy and light chain plasmids transfection following the manufacturer’s instruction. After 5 days, the antibody containing supernatants were collected. Suitable amount of rProtein A Sepharose® Fast Flow beads (Cytiva) was pre-washed and added into supernatants. After overnight incubation at 4 °C, antibody bound protein A beads were collected with Poly-Prep® Chromatography Columns (BIO-RAD). After 3 times wash with DPBS, mAbs were eluted with Fab elution buffer, then neutralized with Tris-HCl. Buffer exchange was performed with Amicon Ultra-4 Centrifugal Filter (MilliporeSigma) to keep mAbs in PBS for following assays. The numbering of mAbs was based on the order of mouse immunization and cloning. Clones 1-4 were mAbs chosen from enriched clones from RBD-his tag protein immunized C57BL/6J mice. Clone 5-11 were mAbs chosen from RBD-his tag protein immunized BALB/c mice.

### Bi-specific antibody generation

Clone 16 (Clone6-KiH-Clone2) bispecific antibody is a human IgG1-like bispecific antibody, generated based on CrossMab-KiH bispecific constructs, including pFUSE2ss-knobLight-hK, pFUSE2ss-knobheavy-hG1, pFUSE2ss-HoleLight-hK, pFUSE2ss-HoleHeavy-hG1. The design and generation of CrossMab-KiH bi- specific constructs was described in the above plasmid constructs parts. The variable region of Clone 6 heavy chain was cloned into pFUSE2ss-knobheavy-hG1vector. The variable region of Clone 6 light chain was cloned into pFUSE2ss-knobLight-hK vector. Clone6-KiH-Clone2 bispecific antibody was expressed *in vitro* in Expi293F^TM^ cells by co-transfecting four plasmids (clone 6 knob heavy chain plasmid, clone 6 knob light chain plasmid, clone 2 hole heavy chain plasmid, and clone 2 hole light chain plasmid) with ExpiFectamine 293 transfection kit (Thermofisher). The expression and antibody purification protocol was similar to recombinant antibody expression described as above. The bi-specific antibody was efficiently purified by using rProtein A Sepharose Fast Flow antibody purification resin (Cytiva, Cat:#17127901).

### Antibody humanization

In order to humanize the antibody, we first determine the six CDR loops from murine variable domains by using the online free program “IGBLAST” (https://www.ncbi.nlm.nih.gov/igblast/). Followed by apply CDR-grafting technique and graft six CDR loops onto human acceptor frameworks. The framework template selection was based on sequence similarity to close human germline sequence, as well as homology to clinically validated germline sequences. Thereafter, we identify Vernier zone residues through CryoEM structure between Clone 2 and trimeric S protein of SARS-CoV-2 from parent antibody (FR residues of Clone 2 within 5Ǻ of trimeric S protein) and substitute the key residues into the human acceptor framework of Clone 13A.

### ELISA

#### ELISA for Anti-serum Titer Determination

The antibody tilters in sera from pre, first and second bleeds were determined using direct coating ELISA. The 384-well ELISA plates (Corning) were coated with 3 μg/mL SARS-CoV-2 RBD-his tag protein (Sino) in PBS at 4°C overnight. After standard washing with PBS-T washing buffer (phosphate-buffered saline containing 0.05% Tween 20), ELSIA plates were blocked with blocking buffer (2% bovine serum albumin dissolved in PBST and filtered) for 1 hr at room temperature. Serial dilutions of pre-immune, first and second immune anti- sera in blocking buffer were added into plates and for 1hr at room temperature. Plates were washed and incubated with relative goat anti-mouse IgG(H+L)/HRP (Thermo Fisher,1:5000) for 1 h at room temperature. Plates were washed and developed using TMB reagents as substrates (Biolegend) following the manufacturer’s recommended protocol. Reaction was stop with stop solution (1M H3PO4) and absorbance at 450nm was recorded by a microplate reader (Perkin Elmer).

#### ELISA for Anti-RBD antibody clones binding

SARS-CoV-2 RBD protein with 6 X Histidine (Sino) was coated at 3 μg/ml in PBS on a 384-well microtiter plate overnight at 4 °C. After standard washing with PBST and blocked with 2% (w/v) solution of BSA in PBST to remove nonspecific binding, purified anti-RBD antibodies were diluted proportionally in PBST+2% BSA and transferred to the washed and blocked microtiter plates. After one hour of incubation at RT, plates were washed, and RBD-his tag protein-bound antibody was detected with goat-anti-human IgG1 (H+L) with horseradish peroxidase (HRP) conjugated (Invitrogen,1:1000) The plates were washed and developed using TMB substrate solution (Biolegend) according to manufacturer’s recommendation and absorbance at 450nm was measured on a microplate reader after the reaction was stopped by stop solution (1M H3PO4).

### Affinity determination via bio-layer interferometry (BLI)

Antibody binding kinetics for anti-spike mAbs were evaluated by BLI on an Octet RED96e instrument (FortéBio) at room temperature. Two types of measurements were performed. (1) HIS1K biosensors (FortéBio) were first loaded with his-tagged SARS-CoV-2 RBD protein to a response of about 1 nm, followed by a 60 s baseline step in the kinetic buffer (PBS, 0.02% Tween, pH7.4). After that, the biosensors were associated with indicated concentrations of the antibodies (from 50 nM to 0.78125 nM with 2-fold dilutions, where the kinetic buffer was served as the negative control) for 200 s, then dissociated in the kinetic buffer for 1000 s. (2) 25ng/ul of Clone13A-IgG1 antibodies were captured on a AHC biosensor (ForteBio). The baseline was recorded for 60 s in a running buffer (PBS, 0.02% Tween-20, and 0.05% BSA, pH 7.4). Afterward, the sensors were subjected to an association phase for 500 s in wells containing RBD-his diluted in the buffer. In the dissociation step, the sensors were immersed in the running buffer for 1000 s. The dissociation constants KD, kinetic constants Kon and Koff, were calculated by using a 1:1 Langmuir binding model with FortéBio data analysis software.

### Affinity measurement by surface plasmon resonance (SPR)

Kinetics binding measurement for anti-spike mAbs in this study was performed using a Biacore T200 instrument (GE Healthcare). The system was flushed with filtered 1xHBS-P+ running buffer (0.01M HEPES, 0.15M NaCl and 0.05%v/v Surfactant P20, pH 7.4) and all steps were performed at 25℃ chip temperature.

#### Kinetics binding measurement on CM5 Chip (Series S Sensor Chip CM5)

For kinetic binding measurements, CM5 chip surface was activated by injecting a solution of EDC/NHS (GE Healthcare). Mouse anti-human IgG (Fc) mAb (25 μg/ml) was immobilized on the sensor chip by amine coupling, followed by deactivation using 1M ethanolamine. Afterward, anti-spike mAbs (0.1 μg/ml) were then flowed over and captured on anti-human IgG (Fc) mAb-coated surface. Subsequently, gradient diluted his-tagged SARS-CoV-2 RBD solutions (1.875 nM-30 nM, two-fold serial dilution) were injected individually in single-cycle kinetic format without regeneration (30 μl/min, association:180s, dissociation:60s). The binding data were double referenced by blank cycle and reference flow cell subtraction. Processed data were fitted by 1:1 interaction model using Biacore T200 Evaluation Software 3.1.

#### Kinetics binding measurement on NTA Chip

For kinetic binding measurements, NTA chip was activated manually by loading a solution of NiCl2. Histidine-labelled SARS-CoV-2 RBD protein (0.075 μg/ml) was then flowed over the chip and captured on nickel-coated surface. Subsequently, gradient diluted anti-spike mAbs solutions (0.9875 nM-15 nM, two-fold serial dilution) were injected individually in single-cycle kinetic format without regeneration (30μl/min, association:240 s, dissociation:90 s). The binding data were double referenced by blank cycle and reference flow cell subtraction. Processed data were fitted by 1:1 interaction model using Biacore T200 Evaluation Software 3.1.

### SARS-CoV-2 pseudovirus reporter and neutralization assays

HIV-1 based SARS-CoV-2 S pseudotyped virions were generated according to a previous study ^71^. Two plasmids are adopted to generate HIV-1 based SARS-CoV-2 S pseudotyped virions. HIV-1 dual reporter vector expressing mCherry and luciferase (NL4-3 mCherry Luciferase, plasmid#44965) was purchased from Addgene. Plasmid expression a C-terminally truncated SARS-CoV-2 S protein (pSARS-CoV-2Δ19) was obtained from Dr Bieniasz’ lab. In order to generate HIV-1 based SARS-CoV-2 S pseudotyped virions, 15x10^6^ 293FT cells were seeded in a 150 mm plates one day before in 20 ml D10 media. The following day, after the cell density reaches 90%, medium was discarded and replaced with 13 mL serum-free Opti-MEM medium. 20 μg NL4-3 mCherry Luciferase reporter plasmids and 15 μg SARS-CoV-2 (pSARS-CoV-2Δ19) plasmids were mixed thoroughly in 225 μl serum-free Opti-MEM medium. Then 100 μl Lipofectamine 2000 (Invitrogen) were diluted in 225 μl serum-free Opti-MEM medium. Then the diluted plasmid mixture and Lipofectamine 2000 were mixed thoroughly and incubated for 10 mins at RT before adding into cells. After 6 hr, the culture medium was changed back to the completed growth medium, 20 mL for one 150 mm plate. At 48 h after transfection, the 20 mL supernatant was harvested and filtered through a 0.45-μm filter, aliquoted and frozen in -80°C.

Parallelly, the three plasmids-based HIV-1 pseudotyped virus system were utilized to generate (HIV- 1/NanoLuc2AEGFP)-SARS-CoV-2 particles and (HIV-1/NanoLuc2AEGFP)-SARS-CoV-2-SA variant particles. The reporter vector, pCCNanoLuc2AEGFP, and HIV-1 structural/regulatory proteins (pHIVNLGagPol) expression plasmid were gifts from Dr Bieniasz’s lab ^71^. Briefly, 293T cells were seeded in 150 mm plates, and transfected with 21 µg pHIVNLGagPol, 21 µg pCCNanoLuc2AEGFP, and 7.5 µg of a SARS-CoV-2 SΔ19 or SARS-CoV-2 SA SΔ19 plasmid utilizing 198 µl PEI. At 48 h after transfection, the 20- ml supernatant was harvested and filtered through a 0.45-μm filter, and concentrated before aliquoted and frozen in -80°C.

The pseudovirus neutralization assays were performed on 293T-hACE2 cell line ^71^. One day before, 293T- hACE2 cells were plated in a 96 well plate, 0.02 x10^6^ cells per well. The following day, serial dilution of monoclonal IgG from 40 μg/mL (4-fold serial dilution using complete growth medium, 55 μL aliquots) were mixed with the same volume of SARS-CoV-2 pseudovirus. The mixture was incubated for 1 hr at 37 °C incubator, supplied with 5% CO2. Then 100 μL of the mixtures were added into 96-well plates with 293T- hACE2 cells. Plates were incubated at 37°C supplied with 5% CO2. 48 hr later, 1 μL D-luciferin reagent (Perkin Elmer, 33.3 mg/ml) was added to each well and incubated for 5 mins. Luciferase activity was measured with using a microplate spectrophotometer (Perkin Elmer). The inhibition rate was calculated by comparing the OD value to relative negative and positive control wells. For the three plasmids-based HIV-1 pseudotyped virus system, 293T cells were collected and the GFP+ cells were analyzed with Attune NxT Acoustic Focusing Cytometer (Thermo Fisher). The 50% inhibitory concentration (IC50) was calculated with a four-parameter logistic regression using GraphPad Prism 8.0 (GraphPad Software Inc.).

### Cell Fusion assay

#### Vectors and plasmids

Plasmid encoding human ACE2 (hACE2) was obtained from Addgene (hACE2; catalog #1786). The hACE2 2.6 kbp ORF was also blunt-cloned into a third generation HIV vector 3’ of CMV promoter and 5’ of an IRES- puro^r^ cassette to generate pHIV-CMV-hACE2-IRES-Puro. It was inserted into a *piggybac* transposon (Matt Wilson of Baylor College of Medicine, along with the transposase plasmid pCMV-*piggybac*) that had been modified to encode a CMV-IRES-*bsd*^r^ cassette; resultant plasmid was named pT-PB-SARS-CoV-2-Spike-IRES-Blasti. This too was inserted into *piggybac* transposon to make pT-PB-SARS-CoV-2-UK Spike-IRES-Blasti.

#### Cell lines

The HOS cells were stably transduced with a third generation HIV vector encoding *tat*, along with eGFP, mRFP, and bleomycin resistance gene; they were maintained in 200-400 μg/mL phleomycin (Invivogen) and were eGFP and mRFP-positive by flow cytometry. hACE2 was subsequently introduced by VSV G-mediated HIV-based transduction using pHIV-CMV-hACE2-IRES-Puro to produce HOS-3734, which cell lines maintained in selection using 10 μg/mL puromycin (Sigma-Aldrich). TZMbl cells (#JC53BL-13) were obtained from the NIH AIDS Reagent Program. TZMbl cells stably expressing wild type S/UK variant S were created by co-transfecting TZMbl cells with pT-PB-SARS-CoV-2- Spike-IRES-Blasti or pT-PB-SARS-CoV- 2-UK Spike-IRES-Blasti, respectively, along with pCMV-*piggybac* and resistant cells selected with 10 μg/mL blasticidin (Invivogen). The control TZMbl cell line not expressing S was generated by co-transfecting pCMV- *piggybac* with pT-pB-IRES-Blasti and selecting for blasticidin-resistant TZMbl cells.

#### Cell fusion inhibition by monoclonal antibodies

Producer cells (TZMbl-wild type Spike/ Tzmbl-UK Spike) and target cells (HOS-3734) were generated as described above. Ten thousand S-expressing cells (TZMbl-wild type Spike/TZMbl-UK Spike) in 100 µL of medium in the absence of blasticidin were seeded in 96 well plates. After 24 h, 70 μL of four-fold serially diluted antibody was added into producer cells and incubated at 37℃ for 1 hour. At that time 10^4^ target cells (HOS-3734) in 50 μL medium were then added to the producer cells, and after another 24 h cells were lysed in 0.1 mL and RLU measured. Data were analyzed with non-linear regression using GraphPad Prism to determine the neutralization curve and the IC50 values calculated.

### *In vitro* neutralization against authentic SARS-CoV-2

SARS-CoV-1 (USA-WA1/2020) was produced in Vero-E6 cells and tittered as described previously ^72^. SARS- CoV-2 neutralization was assessed by measuring cytotoxicity. 5x10^5^ Vero-E6 cells were plated per well of a 96-well plate. The following day, serial dilutions of antibodies were incubated with 2.5x10^3^ plaque forming units (PFU) SARS-CoV-2 for 1 hour at room temperature. SARS-CoV-2 neutralization was assessed by measuring cytotoxicity. 5x10^5^ Vero-E6 cells were plated per well of a 96-well plate. The following day, serial dilutions of antibodies were incubated with 2.5x10^3^ PFU SARS-CoV-2 for 1 hour at room temperature. The medium was then aspirated from the cells and replaced with 100 µl of the antibody/virus mixture. After 72 hours at 37°C, 10 µl of CellTiter- Glo (Promega) was added per well to measure cellular ATP concentrations. Relative luminescence units were detected on Cytation5 (Biotek) plate reader. All conditions were normalized to an uninfected control. Each condition was done in triplicate in each of three independent experiments.

### Focus reduction neutralization test

Serial dilutions of mAbs or sera were incubated with 102 focus-forming units (FFU) of different strains or variants of SARS-CoV-2 for 1 h at 37°C. Antibody-virus complexes were added to Vero-TMPRSS2 cell monolayers in 96-well plates and incubated at 37°C for 1 h. Subsequently, cells were overlaid with 1% (w/v) methylcellulose in MEM supplemented with 2% FBS. Plates were harvested 24 h later by removing overlays and fixed with 4% PFA in PBS for 20 min at room temperature. Plates were washed and sequentially incubated with an oligoclonal pool of SARS2-2, SARS2-11, SARS2-16, SARS2-31, SARS2-38, SARS2-57, and SARS2-71^4 73^. The anti-S antibodies and HRP-conjugated goat anti-mouse IgG (Sigma, 12-349) in PBS supplemented with 0.1% saponin and 0.1% bovine serum albumin. SARS-CoV-2-infected cell foci were visualized using TrueBlue peroxidase substrate (KPL) and quantitated on an ImmunoSpot microanalyzer (Cellular Technologies).

### *In vivo* efficacy testing against authentic SARS-CoV-2

The efficacy of mAbs against replication-competent SARS-CoV-2 virus was evaluated *in vivo*, using both a prophylactic setting where the animals were treated with mAb prior to viral infection, and a therapeutic setting where the animals were treated post infection. These experiments were performed in an animal BSL3 (ABSL3) facility. The replication-competent SARS-CoV-2 (USA-WA1/2020) virus was produced in Vero E6 cells, and the titer was determined by plaque assay using WT VeroE6.

The K18-hACE2 mice (B6.Cg-Tg(K18-ACE2)2Prlmn/J) were purchased from the Jackson Laboratory and bred in house using a trio breeding scheme. Mice were sedated with isoflurane, and infected via intranasal inoculation of 2,000 PFU (20x LD50) SARS-CoV-2 (USA-WA1/2020) virus administered in 50uL of DPBS. Six to eight-week-old K18-hACE2 littermate-controlled mice, mixed gender (male / female) mice were divided randomly into three groups, and administered with 20 mg/kg (of mice body weight) Clone 2, Clone 6 or placebo / control, via intraperitoneal (IP) injection. For prophylactic experiment, the mAb drug / placebo treatment was 24h prior to infection; for therapeutic experiment, the treatment was 18h post-infection. The control for the prophylactic experiment was DPBS, and the control for the therapeutic experiment was isotype control hIgG1, where both controls are similar (no effect on disease progression). Survival, body conditions and weights of mice were monitored daily for 10 consecutive days.

### *In vivo* efficacy testing of humanized Clone13A to authentic SARS-CoV-2 virus

10-12-week-old littermate-controlled female and male K18hAce2Tg+ mice were pretreated with 20 mg/kg of either control hIgG1 (purchased from BioXCell) or clone 13A mAb (produced by the Chen lab) administered IP in 300 uL of DPBS. 24 hours later, mice were anesthetized with isoflurane, and SARS-CoV-2 isolate USA-WA1/2020, or Delta variant (B.1.617.2), was inoculated intranasally at a dose of 2x103 PFU/mouse (determined using wild type Vero E6) in 50 μL of DPBS. Weights were obtained daily for 10 days following infection, and mice were euthanized when morbid.

### Fab generation

The Fab fragments of Clone 2 and Clone 6 were generated from full length IgGs of Clone 2 and Clone 6 using a commercial PierceTM Fab Preparation Kit (Thermo Fisher). All procedures were performed following the manufacturer’s instructions. Briefly, 2 mg the whole IgGs of Clone 2 and Clone 6 were digested with immobilized papain at 37 °C for 4 hr with rotation. Then protein A beads were applied to bind the Fc fragments and undigested IgG. Then Fab fragments were recovered in the flow-through fraction, and further purified by size exclusion chromatography using a Superdex 200 10/300 column (GE Healthcare) in 30 mM Tris pH 8.0, 100 mM NaCl. The mondispersed peak of Fab fragments was pooled and concentrated for subsequent analysis.

### Cryo-EM sample preparation and data collection

The purified SARS-CoV-2 spike trimer at a final concentration of 0.3 mg/mL (after mixture) was mixed with Clone 2 or Clone 6 Fab at a molar ratio of 1:2 at 4 °C for 30mins. Then 3 μl of the protein mixture was applied to a Quantifoil-Cu-2/1-3C grid (Quantifoil) pretreated by glow-discharging at 15 mA for 1 min. The grid was blotted at 4 °C with 100% humidity and plunge-frozen in liquid ethane using FEI Vitrobot Mark IV (Thermo Fisher). The grids were stored in liquid nitrogen until data collection.

Images were acquired on a FEI Titan Krios electron microscope (Thermo Fisher) equipped with a Gatan K3 Summit direct detector in super-resolution mode, at a calibrated magnification of 81,000× with the physical pixel size corresponding to 1.068 Å. Detailed data collection statistics for the Fab-spike trimer complexes are shown in a supplemental table. Automated data collection was performed using SerialEM ^74^.

### Cryo-EM data processing

A total of 2,655 and 1,766 movie series were collected for Clone 2 Fab-S trimer complex and Clone 6 Fab-S trimer complex, respectively. The same data processing procedures were carried out for each complex as described below. Motion correction of the micrographs was carried out using RELION ^75^ and contrast transfer function (CTF) estimation was calculated using CTFFIND4 ^76^. Particles were picked automatically by crYOLO ^77^, followed by 2D and 3D classifications without imposing symmetry. The 3D classes with different S trimer conformations were then processed separately by consensus 3D refinement and CTF refinement. Image processing and 3D reconstruction using cryoSPARC ^78^ produced similar results. For each state of the Clone 6 Fab-S trimer complex, multibody refinements were then carried out in RELION by dividing the complex into individual rigid bodies (3 refinements each with a rigid body containing a unique Fab, RBD, and the N-terminal domain (NTD) of spike S1 subunit, and another rigid body for the rest of the spike ectodomain trimer). For each state of the Clone 2 Fab-S trimer complex, local masked 3D classification without image alignment was performed focusing on one Fab-RBD region and the best class of particles was selected for consensus refinement of the whole complex. Subsequently, multibody refinement was performed as described above for the rigid body containing the focused region. The 3D reconstruction of the other Fab-RBD regions were obtained with the same procedure. The final resolution of each reconstruction was determined based on the Fourier shell correlation (FSC) cutoff at 0.143 between the two half maps ^79^. The final map of each body was corrected for K3 detector modulation and sharpened by a negative B-factor estimated by RELION ^80^, and then merged in Chimera for deposition. The local resolution estimation of each cryo-EM map is calculated by RELION^75^. See also **Extended Data Fig. 7** and **Table 1**.

### Model building and refinement

The structure of the ectodomain of SARS-CoV-2 spike trimer (PDB 6VSB) was used as an initial model and docked into the spike trimer portion of the cryo-EM maps using Chimera ^81^. The initial models of Clone 2 and Clone 6 Fabs were generated by homology modeling using SWISS-MODEL ^82^, and then docked into the Fab portions of the cryo-EM maps using Chimera ^81^. The initial models were subsequently manually rebuilt in COOT ^83^, followed with iterative cycles of refinement in Refmac5 ^84^ and PHENIX ^85^. The final models with good geometry and fit to the map were validated using the comprehensive cryo-EM validation tool implemented in PHENIX ^86^. All structural figures were generated using PyMol (http://www.pymol.org/) and Chimera ^81^.

### Homology modeling of SARS-CoV-2 variants

The structural models of SARS-CoV-2 variants of RBD were generated by SWISS model82 using the wildtype / WA RBD Cryo-EM structure as a template. The generated structures were aligned with the wild-type RBD in complex with Clone 2, Clone 6, and/or other mAbs. The cryo-EM structures and homology models were analyzed in Pymol.

### Replication, randomization, blinding and reagent validations

Replicate experiments have been performed for all key data shown in this study.

Biological or technical replicate samples were randomized where appropriate. In animal experiments, mice were randomized by cage, sex and littermates.

Experiments were not blinded.

Commercial antibodies were validated by the vendors, and re-validated in house as appropriate. Custom antibodies were validated by specific antibody - antigen interaction assays, such as ELISA. Isotype controls were used for antibody validations.

Cell lines were authenticated by original vendors, and re-validated in lab as appropriate. All cell lines tested negative for mycoplasma.

### Data, resources and code availability

All data generated or analyzed during this study are included in this article and its supplementary information files. Specifically, source data and statistics for non-high-throughput experiments are provided in a supplementary table excel file. High-throughput experiment data are provided as processed quantifications in Supplemental Datasets. Genomic sequencing raw data are deposited to Gene Expression Omnibus (GEO) with the accession code (GSE174635). The models of the mAb:Spike complexes have been deposited in the wwPDB with accession codes (3dSpike-Fab6, 7MW2; 2dSpike-Fab6, 7MW3; 1dSpike-Fab6, 7MW4; 2uSpike-Fab2, 7MW5; 3uSpike-Fab2, 7MW6). The cryo-EM maps of the mAb:Spike complexes have been deposited in EMDB with accession codes (3dSpike-Fab6, EMD-24060; 2dSpike-Fab6, EMD-24061; 1dSpike- Fab6, EMD-24062; 2uSpike-Fab2, EMD-24063; 3uSpike-Fab2, EMD-24064), respectively. Codes that support the findings of this research are being deposited to a public repository such as GitHub. Additional information related to this study are available from the corresponding authors upon reasonable request.

## Extended Data Figure Legends

**Extended Data Figure 1.**
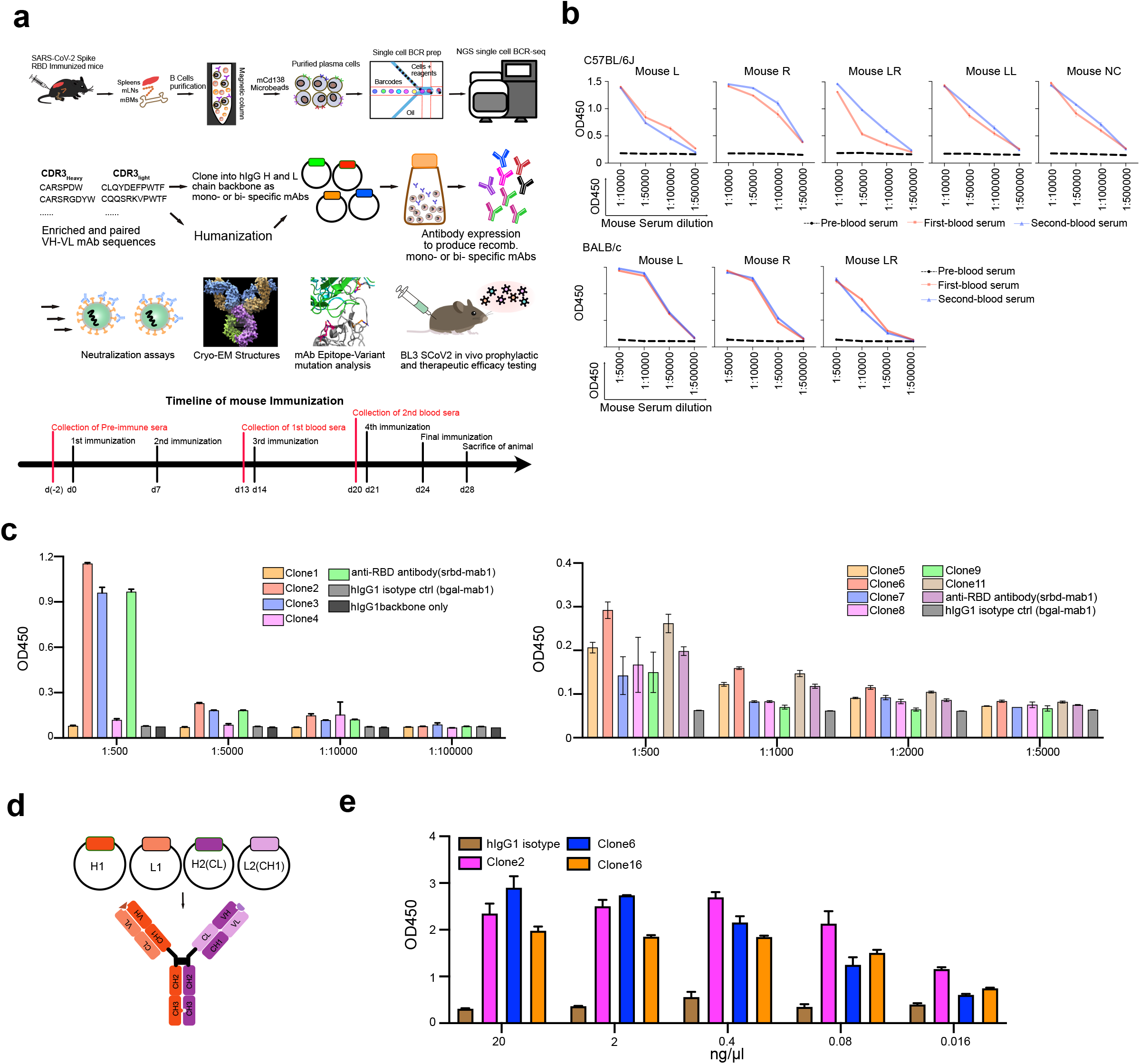
Measurement of mouse serum titers in SARS-CoV-2 Spike RBD immunized mice in this study. **a**, Schematic of experiments. This schematic illustrates neutralizing antibody identification process through RBD-his tag protein mouse immunization-single B cell sequencing (**Top**), along with main assays of downstream analyses (**Middle**). The paired heavy chain and light chain sequences of the B cells were obtained using 10X Genomics VDJ sequencing. Antibodies were reconstructed by cloning of IgG heavy and light chains into human IgG1 backbone and expressed as recombinant monospecific or bispecific mAbs. Lead antibody clones were subjected to characterizations including neutralization assays, BL3 level anti- authentic SARS28 CoV-2 efficacy testing, and structural analyses by Cryo-EM. (Bottom) A timeline of mouse immunization for antibody development. **b**, SARS-CoV-2 RBD reactivity ELISA result of serum samples from different RBD immunized C57BL/6J (b) and BALB/c (c) mice. **c**, SARS-CoV-2 RBD reactivity ELISA result of recombinant monospecific mAb clones identified from single BCR sequencing of RBD immunized C57BL/6J (top) and BALB/c (bottom) mice. **d**, Schematics of construct design and antibody structure of bispecific antibodies used in this study. **e**, SARS-CoV-2 RBD reactivity ELISA result of top monospecific mAb clones (Clones 2 and 6) and a bispecific mAb clone (Clone 16). Source data and additional statistics for experiments are provided in a supplemental excel file.

**Extended Data Figure 2.**
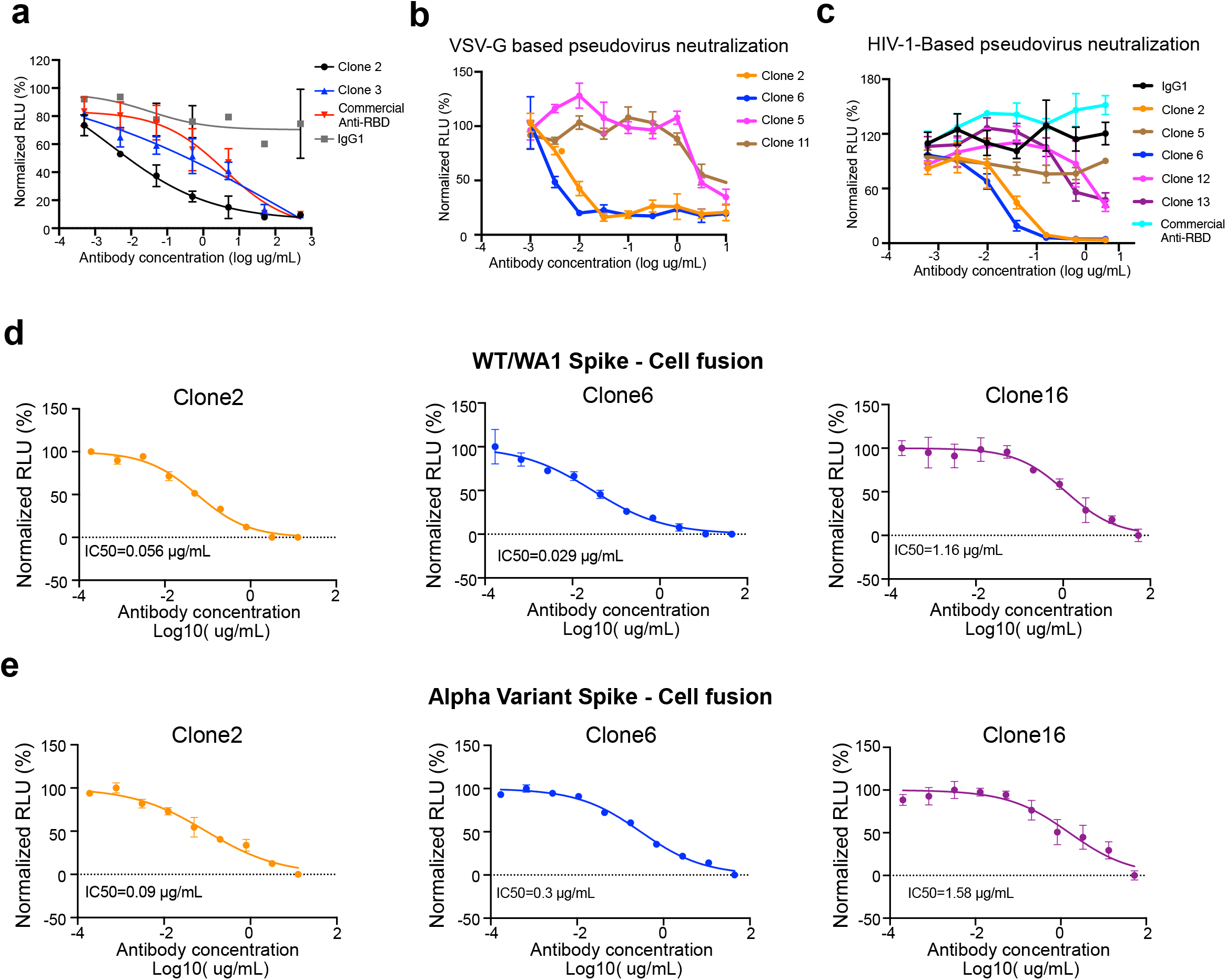
Neutralization capability testing of antibody clones with HIV1-based and VSV-G-based WT SARS-CoV-2 spike pseudotyped virus. **a**, Neutralization assay on Clone 2, and Clone 3 using WT SARS-CoV-2 Spike pseudotyped lentivirus carrying a luciferase reporter. **b**, Neutralization assay on Clone 2, Clone 6, Clone 5 and Clone 11 using WT SARS-CoV-2 Spike pseudotyped VSV- virus carrying a luciferase reporter. **c**, Neutralization assay on Clone 2, Clone 5, Clone 6, Clone 12 and Clone 13, using WT SARS-CoV-2 Spike pseudotyped HIV-lentivirus carrying a luciferase reporter. **d**, Cell fusion assay with SARS-CoV-2 Spike on top mAb clones. **e**, Cell fusion assay with SARS-CoV-2 UK variant Spike on top mAb clones Source data and additional statistics for experiments are provided in a supplemental excel file.

**Extended Data Figure 3.**
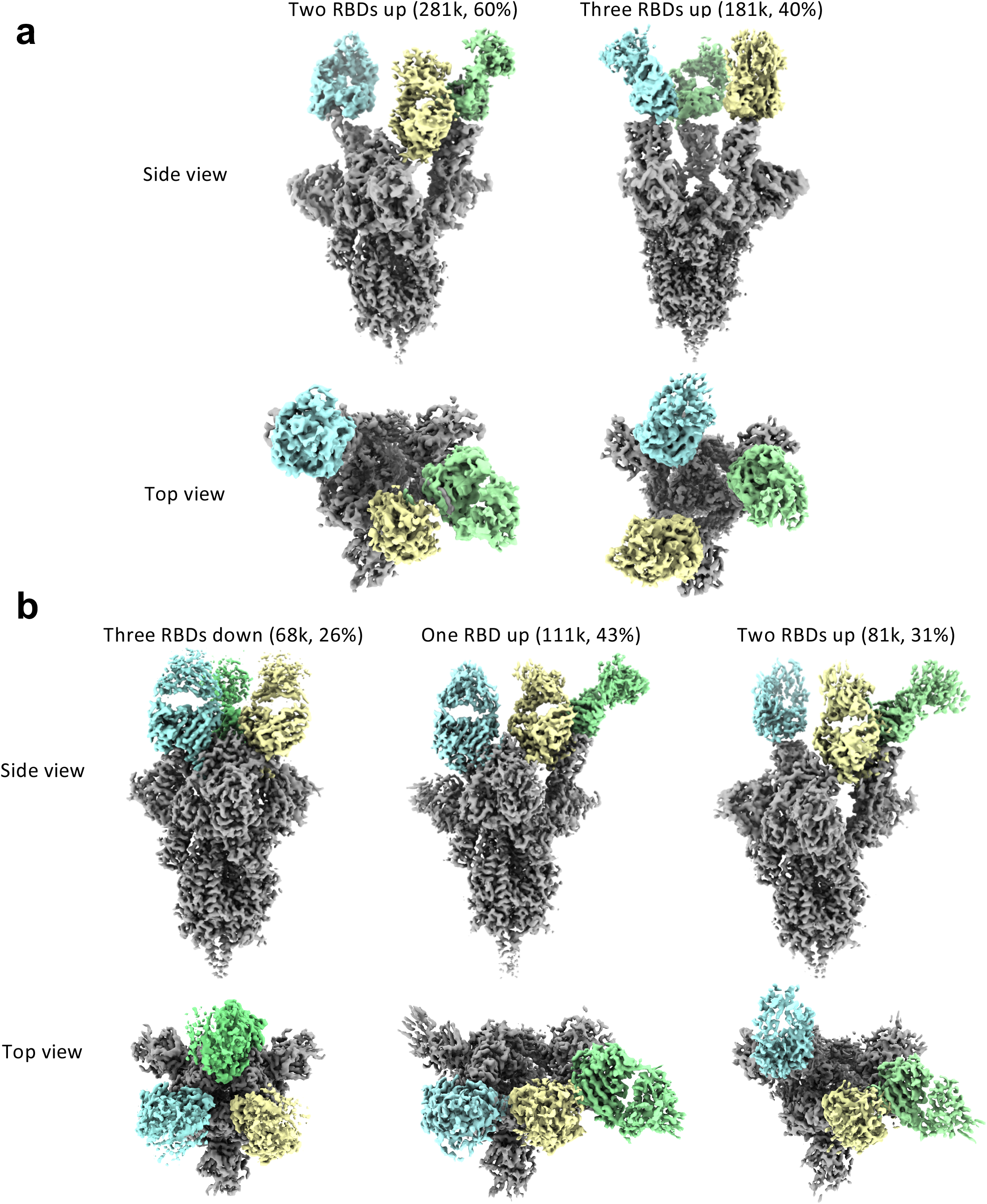
Cryo-EM density maps of the ectodomain of SARS-CoV-2 S trimer (gray) in complexes with Clone 2 (a) or Clone 6 (b) Fabs (green, yellow and cyan).

**Extended Data Figure 4.**
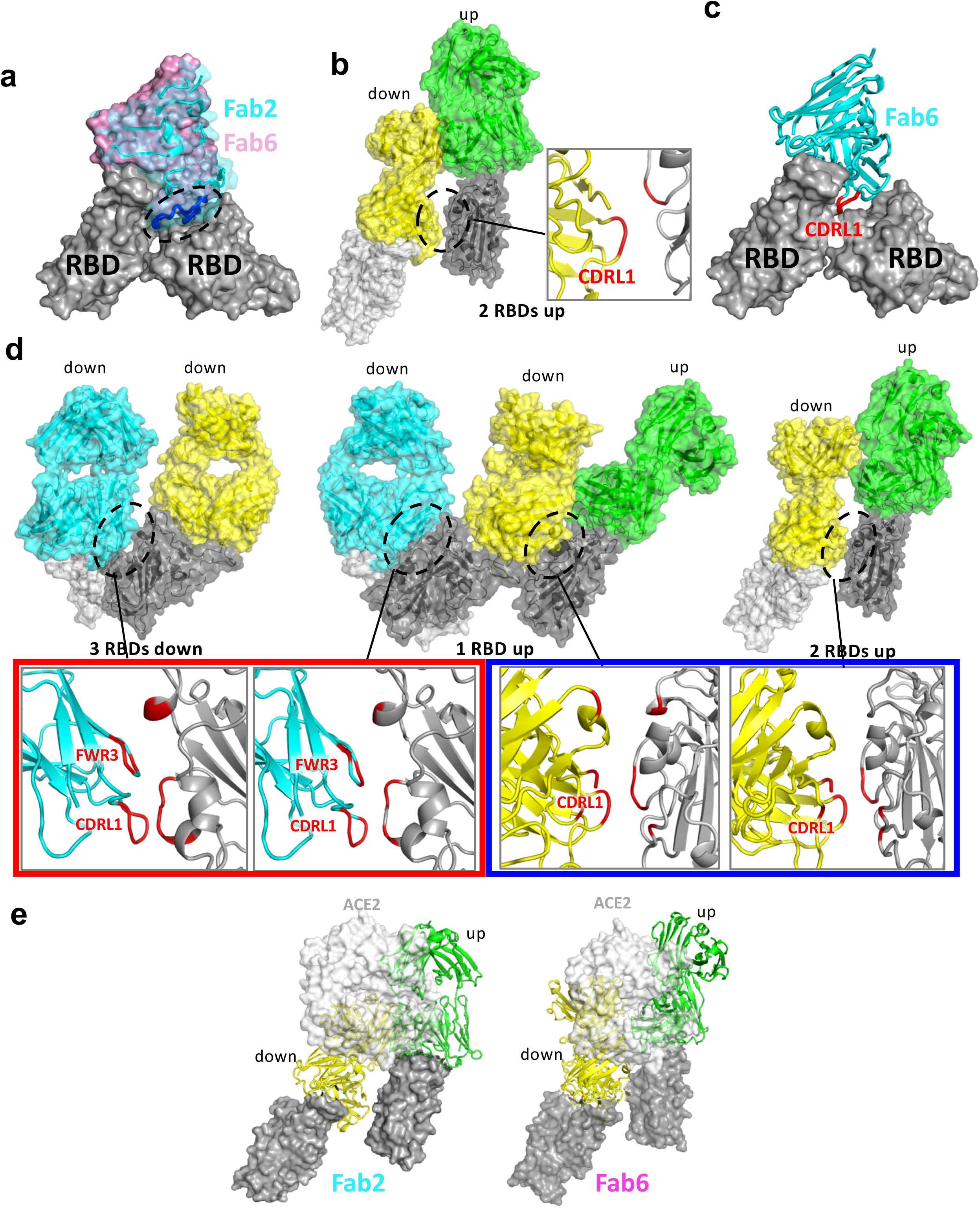
Additional binding interfaces between spike RBD and both clone Fabs. **a,** Overlay of a down-RBD-bound Clone 2 Fab (cyan ribbons with transparent surface) onto a down-RBD-bound Clone 6 Fab (cyan surface) reveals a steric clash between the Clone 2 Fab and a neighboring RBD. **b**, The additional binding interface between a down-RBD-binding Clone 2 Fab and an adjacent up-RBD in S trimers with 2 RBDs up. **c**, The down-RBD-binding Clone 6 Fab sits on top of two adjacent down-RBDs in S trimers with 2 or 3 RBDs down. The CDRL1 loop highlighted in red inserts between two adjacent RBDs. **d**, The additional binding interfaces between a down-RBD-binding Fab and adjacent down- or up-RBD in all Clone 6 Fab-S trimer complexes. The residues involved in the interactions are highlighted in red. **e**, ACE2 bound to an up-RBD has additional steric clashes with a neighboring down-RBD bound Clone 2 (left) or Clone 6 (right) Fab in spike trimers with either 1 RBD up or 2 RBDs up. ACE2 is shown as light gray surface, Clone 2 and 6 Fabs are shown as yellow (on down-RBD) or green (on up-RBD) ribbons, and RBDs are shown as gray surfaces.

**Extended Data Figure 5.**
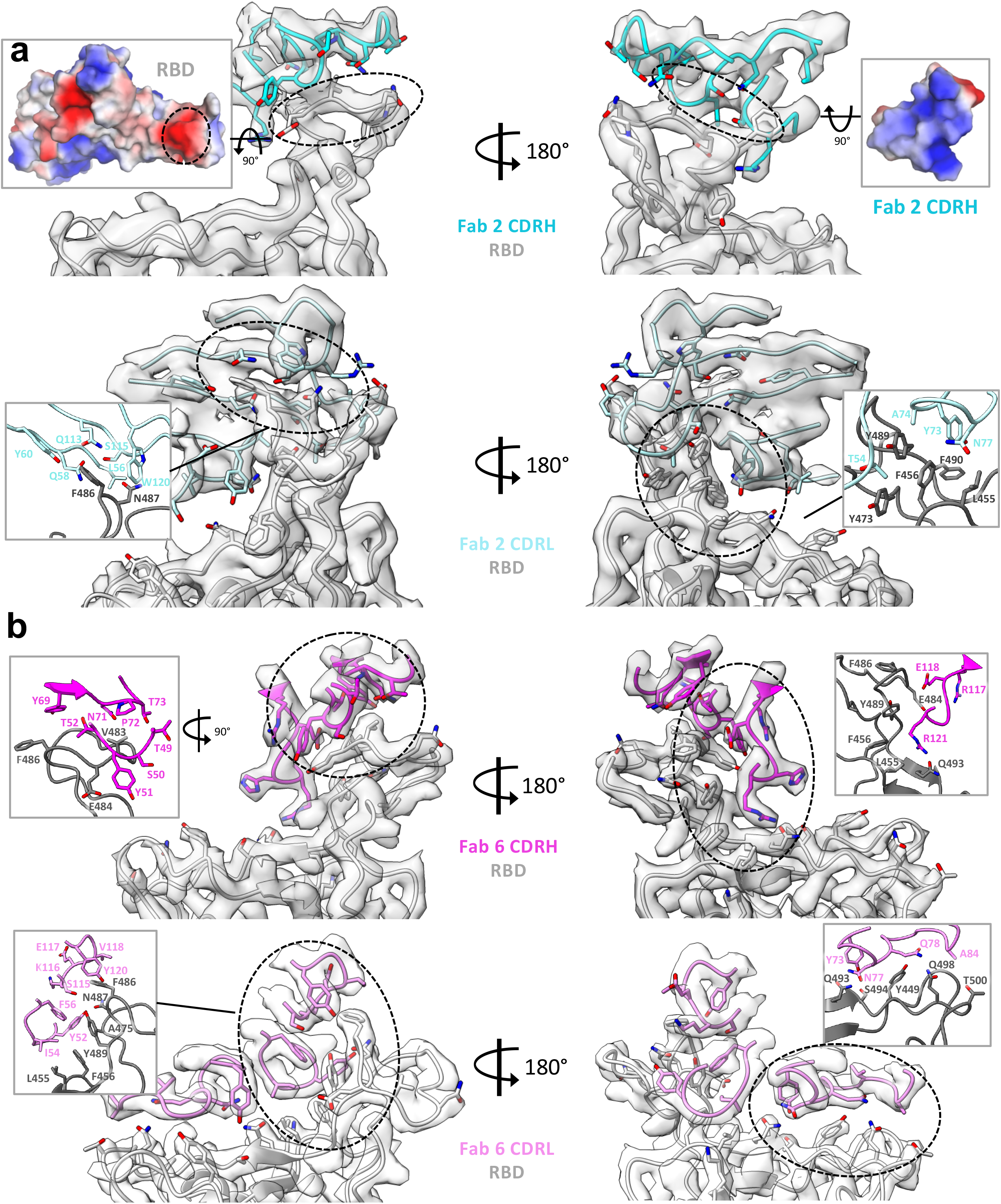
Detailed atomic interactions at the spike RBD-Fab interfaces. **a**, Cryo-EM maps of the spike RBD binding interfaces with the Clone 2 Fab CDRH loops (upper panel) and CDRL loops (lower panel), with the fitted models in ribbon representation. The key interactions and electrostatic surfaces (blue, positively charged; red, negatively charged) at the binding interfaces are shown in respective insets. **b,** Cryo-EM maps of the spike RBD binding interfaces with the Clone 6 Fab CDRH loops (upper panel) and CDRL loops (lower panel), with the fitted models in ribbon representation. The key interactions are shown in respective insets.

**Extended Data Figure 6.**
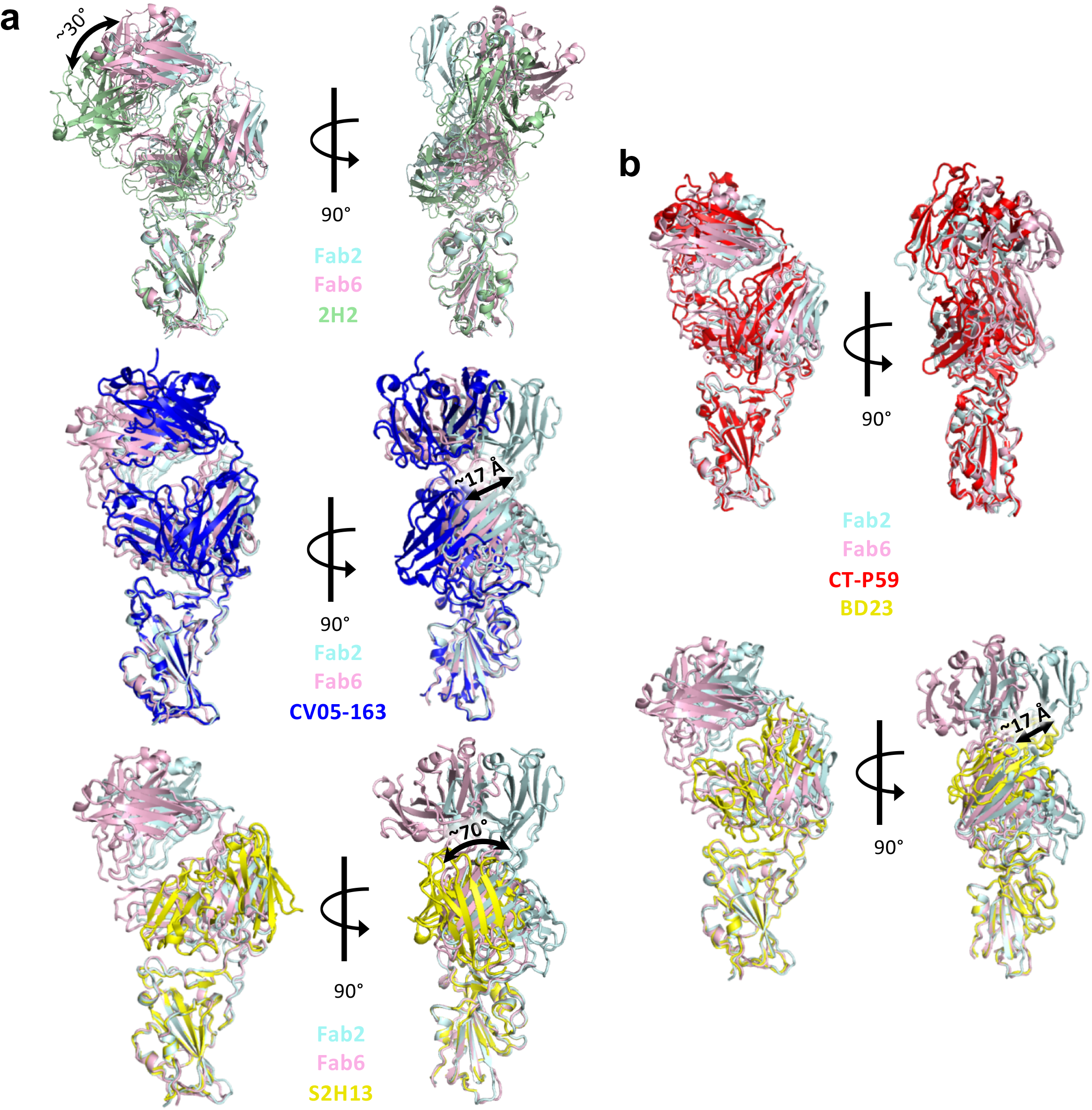
Comparison of binding orientations between Clone 2/6 and previous reported antibodies on SARS-CoV-2 spike RBD, with the RBD portions overlaid. **a**, Three previously reported antibodies bind the spike RBD in overall similar orientations as those of Clone 2/6, but with substantial rotations or shifts. **b**, Two previous reported antibodies resemble the binding conformations of Clone 2/6 on spike RBD, however with the positions of heavy chains and light chains swapped. The PDB IDs of the published Fab-RBD structures: 2H2, 7DK5; CV05-163, 7LOP; S2H13, 7JV4; CT-P59, 7CM4; BD23, 7BYR.

**Extended Data Figure 7.**
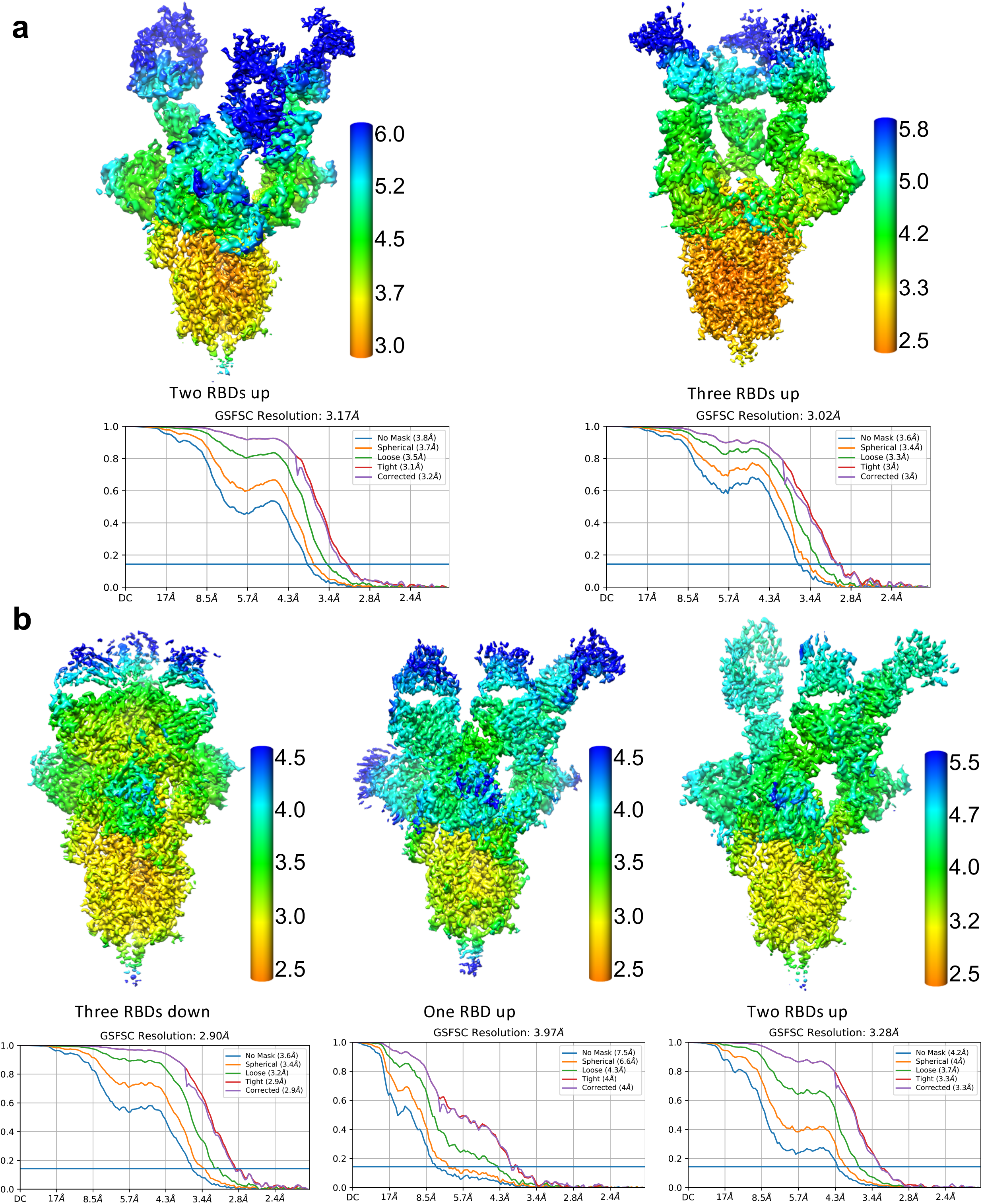
Local resolution estimations of the cryo-EM maps of SARS-CoV-2 S ectodomain trimer (gray) in complexes with Clone 2 (a) or Clone 6 (b) Fabs. Fourier shell correlation (FSC) curves of the half-maps of each complex structure from gold standard refinement calculated by cryoSPARC are also shown, respectively.

**Extended Data Figure 8.**
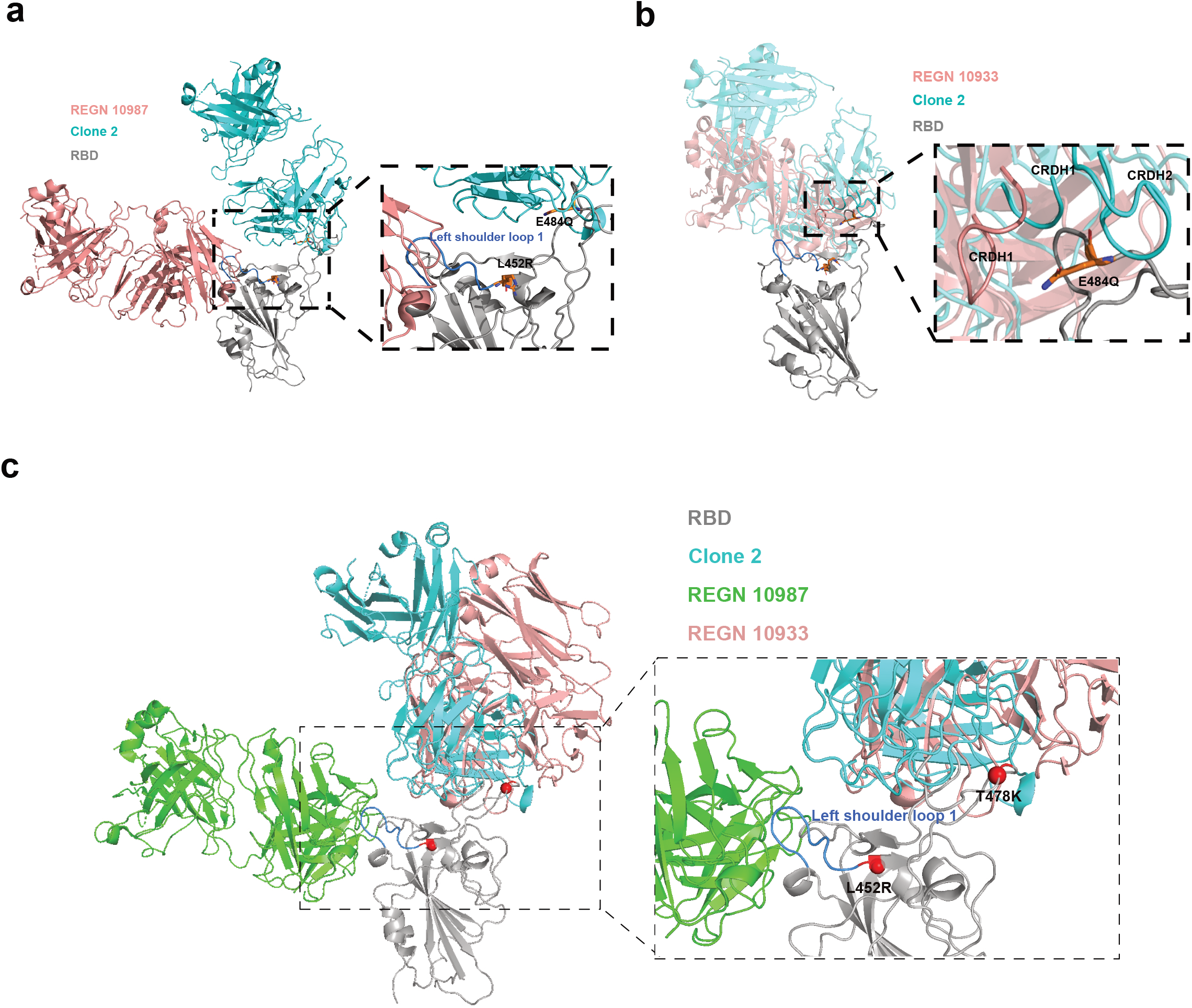
Epitope and mutation analysis of the Fab-spike RBD interfaces for lead mAb clone in comparison with representative existing clinical antibodies. Structural comparison of Clone 2, REGN 10933 and REGN 10987 revealed their distinct RBD epitopes and varied susceptibility to mutations of Delta variant. RBD structural model of Delta in complex with REGN 10987 (A, Pink, PDB: 6XDG), REGN 10933 (B, Pink, PDB: 6XDG) and Clone 2 (Cyan) is shown. **a**, The epitope of REGN 10987 are mainly distributed in the left shoulder loop 1 (blue) region, which extends to L452R mutation site, while the clone 2 mainly targets the right ridge of RBD. **b**, The paratopes of REGN 10933 and clones targeting the E484Q of RBD are labeled and shown. The remaining parts of the antibody structures were set 50% transparent. **c**, Overlay of structures of REGN 10933, RGEN 10987 and Clone 2 Fab with RBD, with analysis of representative key residues.

## Supplemental Datasets

**Dataset S1 Single cell BCR sequencing processed data**

**Supplemental Source Data and Statistics**

Original data and statistics for non-high-throughput data used in figure generation.

**“CoVmAbBis_Statistics_Original_data_v7.xlsx”**

